# Conserved roles of GATA4 and its target gene TBX2 in regulation of human cardiogenesis

**DOI:** 10.1101/2025.07.11.664341

**Authors:** Nicola Graham, Pavel Kirilenko, Ilya Patrushev, Ewan D. Fowler, Peter Kille, Michael Gilchrist, Nick D.L. Owens, Branko Latinkic

**Affiliations:** School of Biosciences, Cardiff University, Cardiff, United Kingdom; The Francis Crick Institute, London, United Kingdom; Department of Clinical and Biomedical Sciences, Exeter University, United Kingdom

## Abstract

The transcription factor (TF) GATA4 is a key mediator of cardiogenesis. GATA4 regulates cardiogenesis through the expression of its target genes, only some of which have been identified. We have used a gain of function model based on pluripotent embryonic ectoderm explants from *Xenopus* embryos expressing GATA4, to identify a set of downstream targets of GATA4 which are also regulated by Nodal, a known cardiogenic signal. GATA4 was shown to be required for the expression of target genes *tbx2* and *prdm1* in vivo, likely acting in a direct fashion by interacting with their regulatory regions. In addition, *tbx2* and *prdm1* are shown to have roles of their own in vivo, as downregulation of *tbx2*, a positive target, and overexpression of *prdm1*, a negative target, interferes with cardiac development in *Xenopus* embryos.

The conservation of the *GATA4-TBX2-PRDM1* regulatory relationship was shown in human iPSC-derived cardiomyocytes. Loss of function of GATA4 lead to downregulation of *TBX2*, upregulation of *PRDM1* expression and failure of cardiogenesis. GATA4-deficient cells failed to form normal cardiomyocytes, with most cells adopting alternative fates and only a small minority expressing an aberrant cardiomyocyte phenotype. Genome-wide transcriptomic analysis documented severe reduction of cardiomyocyte and endothelial cell transcriptomes and upregulation of transcriptional profiles of smooth muscle cells and fibroblasts. Disruption of TBX2 function did not alter cardiomyocyte differentiation efficiency but led to the formation of hypertrophic cardiomyocytes characterised by defective sarcomeres and deficient calcium signalling. In addition, we show that whilst *PRDM1* is not essential for formation of cardiomyocytes it is implicated in suppression of alternative cell fates.

The results presented establish a conserved regulatory relationship between GATA4 and its target genes *TBX2* and *PRDM1* and roles for these genes in the modulation of cardiomyocyte development, expanding the cardiac gene regulatory network and providing further insight into how cardiogenesis proceeds.

## Introduction

Heart development and homeostasis is controlled by the collective action of numerous transcription factors (TFs) which come together to form complex and dynamic gene regulatory networks (GRN) that define and maintain cellular identity. Disruption of the cardiac GRN is associated with the development of congenital heart defects (CHDs), and cardiovascular disease (CVD) in later life (Fahed et al., 2014; Tan et al., 2002). The established core network of TFs that orchestrate cardiac development, includes GATA4/5/ 6, NKX2-5, HAND1, 2, TBX2/ 5/ 20 and proteins belonging to the MADS domain family such as MEF2 and SRF (Akerberg and Pu, 2020; Bruneau, 2013; Olson, 2006; Paige et al., 2015). Evidence from mouse knockout models has shown that the absence of any one of these factors results in an array of severe cardiac defects that almost invariably lead to embryonic lethality (Bruneau et al., 2001; Firulli et al., 1998; Koutsourakis et al., 1999; Kuo et al., 1997; Srivastava et al., 1997; Stennard et al., 2005; Watt et al., 2004) highlighting vulnerability of the cardiac GRN to perturbation.

GATA4 is a core member of the cardiac GRN with well documented roles in all stages of heart development as well as in post-natal life. Early experiments in mice identified an indirect role for GATA4 in visceral endoderm which is essential for cardiac morphogenesis in the overlying mesoderm (Kuo et al., 1997; Molkentin et al., 1997). Subsequent studies have identified cell autonomous roles for GATA4 in the developing heart (Watt et al., 2004; Zeisberg et al., 2005). Evidence for the importance of GATA4 in human heart development is exemplified by the characterisation of a heterozygous mutation, G296S, linked with congenital heart disease (CHD) (Garg et al., 2003). Cardiomyocytes generated from CHD patient-derived induced pluripotent stem cells (iPSCs) show defects in contractility and metabolism (Ang et al., 2016). The molecular basis for this phenotype was shown to be the inability of the mutant GATA4 protein to recruit TBX5 to common targets in chromatin, leading to downregulation of cardiac targets and to aberrant de-repression of endothelial targets (Ang et al., 2016). It is worth noting that this heterozygous mutation in patients is associated with the occurrence of a range of atrial and ventricular septal defects of a severity often requiring surgical correction (Garg et al., 2003). In contrast, heterozygous mice carrying the corresponding mutation appear grossly normal, with a proportion showing mild defects in post-natal atrial septal closure (Misra et al., 2012), highlighting the limitations of mouse models of CHD.

In addition to being required for heart development, GATA4 has been shown to be sufficient to drive cardiomyocyte differentiation of permissive embryonic cells either on its own in pluripotent ectodermal explants from *Xenopus* embryos (Latinkić et al., 2003), or together with Baf60c in mouse early embryonic mesoderm (Takeuchi and Bruneau, 2009). Furthermore, GATA4 is used in most cocktails of cardiac TFs for reprogramming of non-cardiac cells to cardiomyocytes (Garry et al., 2022). For example, *GATA4* along with additional cardiac GRN members *MEF2* and *TBX5* is sufficient for reprogramming of fibroblasts to cardiomyocytes (Garry et al., 2022; Xie et al., 2022). The requirement for GATA4 for normal cardiac development, as outlined above, and the ability of GATA4 to steer the genomic program of non-cardiomyocytes towards a cardiomyocyte fate reaffirms it as a key component of the cardiac GRN.

Mechanistic studies focused on mapping of GATA4 occupancy sites in chromatin in different models such as mouse embryos, cardiomyocytes derived from human iPSCs and adult hypertrophic hearts, are beginning to identify the roles of GATA4 at the chromatin level by defining its stage-, signal- and cell type-specific targets. Besides the core cardiac GRN, numerous additional transcription factors regulate multiple aspects of cardiac development such as cardiac cell type diversification and morphogenesis. The emerging expanded cardiac GRN includes genes such as *TBX2* which has been shown to regulate the formation of the Atrio-Ventricular Canal (AVC) by repressing the transcriptional program for chamber cardiomyocytes. To describe an expanded cardiac GRN, it will be necessary to obtain and integrate a better knowledge of regulatory relationships between its nodes.

A simple model for studying the cardiac GRN is offered by overexpression of GATA4 in pluripotent ectodermal explants from blastula stage *Xenopus* embryos. GATA4 is sufficient to reprogram explant cells fated to become epidermis towards cardiomyocytes in a process which mimics normal heart development by producing beating cardiac tissue (Latinkić et al., 2003). In this model, in addition to cardiomyocytes GATA4 induces endothelial cells as well as anterior endoderm, in keeping with its known roles in vivo. GATA4-driven cardiogenesis was shown to be enhanced by Wnt antagonism (Latinkić et al., 2003), in agreement with the well documented role of Wnt antagonism in heart development.

Besides GATA4 expression, pluripotent explants from *Xenopus* embryos can be directed towards cardiomyocyte fate by Nodal/Activin signalling, the principal inducer of mesoderm and endoderm in vertebrate embryos (Logan and Mohun, 1993; Takahashi et al., 2000). In *Xenopus* explants, Nodal signalling acts in a concentration dependent manner, with a high dose required for induction of endoderm and cardiomyocytes and a lower dose for skeletal muscle (Logan and Mohun, 1993; Takahashi et al., 2000).

As both GATA4 and Nodal induce cardiomyocytes, we used both cardiogenic triggers in this report to identify and characterise putative components of the cardiac GRNs as co-regulated differentially expressed genes (DEGs). A subset of DEGs were validated as GATA4 targets in vivo as well as in explants. Interference with normal expression of two such targets, *tbx2* and *prdm1* is shown to affect heart development in *Xenopus* embryos. The wider relevance of the data obtained in *Xenopus* is shown by our results which demonstrate the essential roles of GATA4 and its target gene *TBX2* in differentiation of human cardiomyocytes from iPS cells.

## Results

### Identification of GATA4 targets under cardiogenic conditions

To dissect the genomic program for cardiogenesis regulated by GATA4 we employed a well-established gain of function model using pluripotent ectodermal explants from blastula stage *Xenopus* embryos (Latinkić et al., 2003). As GATA4 plays key roles in the development of multiple cell types of mesodermal and endodermal origin, and is sufficient to specify endodermal, endothelial and blood cells in *Xenopus* animal cap explants, we designed an experimental approach for selection of cardiogenesis-related genes regulated by GATA4. This consisted of defining a set of genes regulated by cardiogenic conditions: GATA4 (G4), GATA4 in the presence of Wnt antagonist Dkk-1 (G4D), which enhances cardiogenesis, and high level of Nodal/Activin signalling, known to induce cardiogenesis (Fig. 1, S1). At the same time, these genes were not regulated by a lower level of Nodal/Activin which induces skeletal but not cardiac muscle, and by Dkk-1 alone (Fig. 1, S1). In this model it takes approximately 2 days for GATA4 to induce cardiac differentiation as assessed by expression of cardiomyocyte markers (Fig. 1A, S1A).

**Figure 1.**
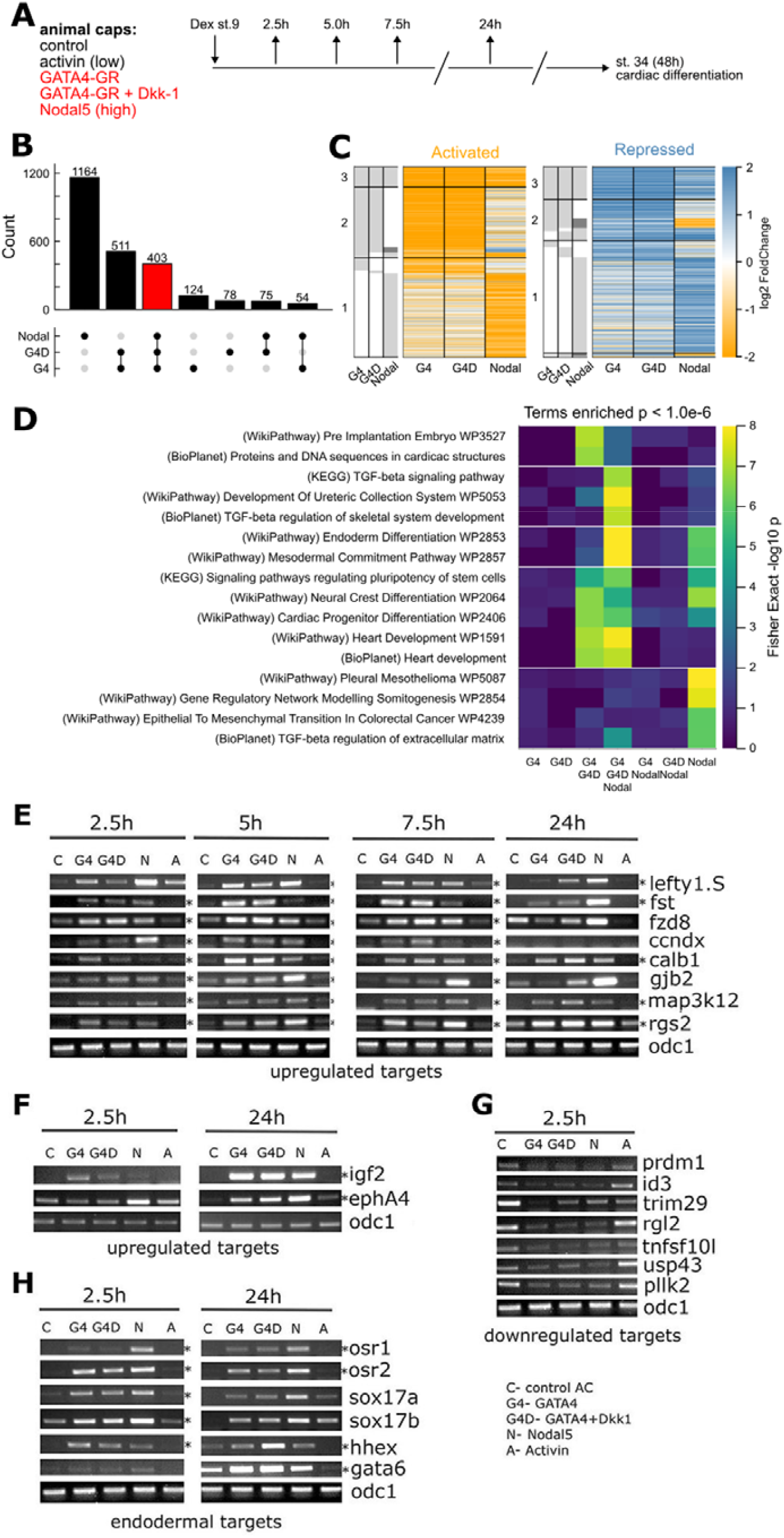
Identification and validation of cardiogenesis-related targets of GATA4. (A) Experimental outline. Animal cap explants from blastula stage embryos were excised and incubated for the indicated time before being harvested and subjected to RNA-seq analysis. The samples injected with mRNAs coding for proteins known to induce cardiac tissue are indicated in red. GATA4-GR was activated by incubation in 1 mM dexamethasone. The control sample were uninjected animal cap explants. Animal caps treated with a low dose of soluble activin, known to induce skeletal but not cardiac muscle (Fig. S1), was used as an additional control and a method to select genes responsive to cardiac inducing conditions. (B) Upset plot describing intersection of differentially expressed genes irrespective of timepoint compared to control. Bars indicate total number of genes differentially expressed in combination of conditions indicated in lower plot at any timepoint. (C) Heatmap showing genes activated and repressed per condition irrespective of timepoint. Left grey heatmap indicates whether gene was significantly differentially expressed in indicated condition. Corresponding heatmap divided per time point shown in Fig. S1E. (D) Gene set enrichment calculated using Enrichr. Heatmap shows -log10 FDR for enrichment of indicated term for genesets defined by upset plot in (B), within WikiPathway, BioPlanet and KEGG annotations. Heatmap divided into five classes of differing enrichment responses. Geneset enrichments for Gene Ontology is provided in Fig. S2, and table of all tested terms provided in Supplemental Table 1. (E, F) RT-PCR validation of a subset of upregulated DEGs. Time points where each DEG was validated are indicated by an asterisk. o*dc1* expression is used as a loading control. The same *odc1* data is shown in all panels where the same cDNA samples were used. (G) RT-PCR validation of downregulated DEGs at 2.5h of incubation. All DEGs were validated as downregulated by GATA4-GR, GATA4-GR + Dkk-1 and Nodal 5 compared to the level of expression in uninjected or activin treated animal caps. (H) RT-PCR validation of DEGs linked with endoderm development at 2.5h and 24h. Time points where each DEG was validated are indicated by an asterisk.

We focussed on the early stages of the genomic program which was expected to include immediate-early and delayed-early targets, by sampling at 2.5h, 5h and 7.5h after excision of animal cap explants and activation of inducible GATA4-GR fusion by addition of dexamethasone (DEX). We performed polyA+ RNA-seq and calculated DEGs at each stage relative to control. The three conditions produced a highly concordant impact on gene expression for each time point (Fig. S1B-D); therefore, we combined these to define DEGs per condition. We found 1,092 DEGs (690 activated, 402 repressed) for GATA4 (G4), 1,067 (602 activated and 465 repressed) for GATA4 + Dkk-1 (G4D), and 1,696 (829 activated and 867 repressed) for Nodal. Comparing between conditions we found exceptional overlap of 914 genes differentially expressed in G4 and G4D, 83.7% and 85.7% of their genes respectively (Fig. 1B). The overlap with Nodal DEGs was more limited with 403 genes shared between the three cardiogenic treatment groups (Fig. 1B). We performed gene set enrichment using Enrichr (Kuleshov et al., 2016) (Fig. 1D, S2; Supplemental Table 1), and we calculated enrichments for all intersection groups indicated in the upset plot (Fig. 1B). We found a clear induction of cardiac gene expression programmes in the intersection of GATA4 and Nodal DEGs, with terms such as Heart Development (FDR < 10^-7^, intersection of G4/G4D/Nodal) and Cardiac Progenitor Differentiation (FDR < 10^-5^, intersection of G4/G4D/Nodal). We further found an enrichment in Gene Ontology (GO) terms related to transcription factor driven regulation of gene expression in the intersection of GATA4 and Nodal conditions, with enrichment of terms such as Regulation Of Transcription By RNA Polymerase II (FDR < 10^-23^) (Fig. S2).

Validation confirmed DEGs which satisfied the selection criteria (regulated by G4, G4D and high Nodal but not low Nodal/Activin) at all time points such as *fst, map3k12, rgs2* (Fig. 1E). Other DEGs were validated at some but not all time points.

Upregulated DEGs include known Nodal target *lefty1* (Cheng et al., 2000); at 2.5h *lefty1*.*S* is strongly induced by Nodal, as expected from a direct target of Nodal signalling (Fig. 1E). However, as the gene is also induced by low level of activin/Nodal signalling at that time, it is not validated as a DEG induced by cardiogenic conditions. The expression of *lefty1*.*S* induced by cardiogenic but not non-cardiogenic conditions is sustained beyond 2.5h, thus identifying it as a DEG (Fig. 1E). Additional upregulated DEGs include *igf2* and *ephA4* (Fig. 1F), Tbx family transcription factors *tbx2* and *tbx5* (Fig. 2D) and *gata2, hoxc13* and *lmo2* (Fig. S3).

**Figure 2.**
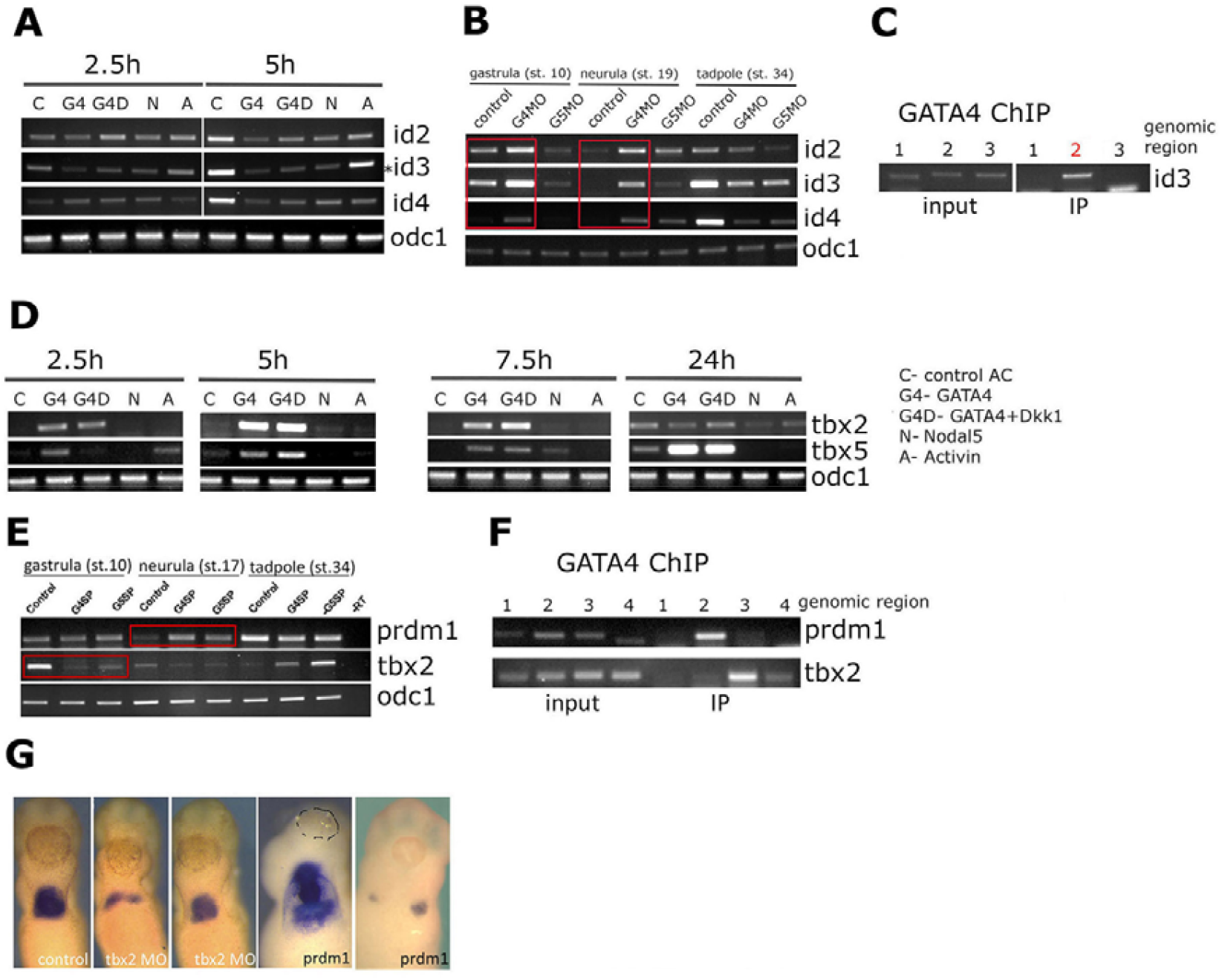
*id3, tbx2* and *prdm1* are regulated by GATA4. (A) RT-PCR analysis for the expression of *id2-4* genes in control (uninjected), GATA4 injected, GATA4+Dkk-1 injected, Nodal 5 injected and activin treated animal cap explants at 2.5 and 5h after excision of animal cap explants and activation of inducible GATA4-GR by 1 mM dexamethasone. *id3* is downregulated by cardiogenic conditions at 2.5h and more strongly at 5h. This is in contrast to family members *id2* and *id4*, which were not picked up in the screen. *odc1* expression is used as loading control. (B) GATA4 is required for regulation of *id2-4* gene expression in vivo. Embryos injected with control Morpholino Oligonucleotides, GATA4 Morpholino Oligonucleotides (G4MO) or GATA5 Morpholino Oligonucleotides (G5MO) were collected at indicated stages and analysed for expression of *id2-4* and *odc1* by RT-PCR. Downregulation of GATA4 (G4MO samples) leads to upregulation of *id2-4* expression at st. 10 and st. 19 (indicated by red box), as assessed by RT-PCR. (C) GATA4 associates with *id3* genomic locus. 3 genomic regions containing GATA sites were amplified from genomic DNA (input) or from DNA immunoprecipitated (IP) by antibodies against HA-tagged GATA4. Region 2 but not 1 and 3 associate with GATA4. (D) Validation of *tbx2* and *tbx5* as GATA4 targets in animal cap explants. Samples collected at indicated time points were analysed by RT-PCR for expression of *tbx2, tbx5* and *odc1. tbx2* is strongly induced by GATA4 and GATA4+Dkk-1 and weakly by Nodal 5 at 5h; *tbx5* is induced by all 3 cardiogenic conditions at 7.5h and by GATA4 and GATA4+Dkk-1 at 5h and 24h. (E) GATA4 is required for regulation of *prdm1* and *tbx2* in vivo. *prdm1* expression in both GATA4 splice-blocking MO (G4SP) and GATA5 splice-blocking MO (G5SP) is upregulated compared to control MO-injected embryos at st.17, whereas *tbx2* expression is downregulated upon GATA4 and GATA5 knockdown (indicated by red boxes). (F) GATA4 binds discrete genomic regions of *prdm1* and *tbx2*. Two additional GATA4-binding regions were identified for *prdm1* and are shown in Fig. S3B. The region 2 is the same as the amplicon 12 in Fig. S3B. Sequences of all positive amplicons are shown in Fig. S3C. (G) Downregulation of *tbx2* and gain of function of *prdm1* lead to heart abnormalities in vivo. Ventral views of representative st. 34 embryos (anterior to the top) with the heart revealed by *myl7* WMISH. Injection of *tbx2* MO leads to cardia bifida or smaller heart region. Injection of *prdm1* mRNA causes defective development of the heart (linear heart tube or cardia bifida). Additional examples and phenotype classification are shown in Fig. S3D,E. Dashed line outlines cement gland in a sample where it is not clear.

In addition to upregulated DEGs, we have identified DEGs which are downregulated in response to cardiac inducing conditions (Fig. 1G). They were most consistently observed at 2.5h, the proximal time point at which direct action of G4, G4D and Nodal is most likely. Downregulated DEGs include transcription factors *prdm1* and *id3*, signalling molecules such as *pllk2, tnfsf10l, rgl2*, E3 ubiquitin ligase *trim29* and ubiquitin peptidase *usp43* (Fig. 1G). Since *id3* is a member of a small family of transcription factors (*id2-4* in *Xenopus*) with described developmental roles including the heart (Cunningham et al., 2017), we have also examined the expression of *id2* and *id4* under the conditions of our screen. As shown in Fig. 2A, in contrast to *id3, id2* and *id4* are not found to be differentially expressed.

Whilst the screen conditions induced a programme of cardiogenesis, none of the three treatments is strictly cardiogenic. Indeed, our GO analysis has indicated enrichment of terms such as Endoderm Differentiation (Fig. 1D) and we found early endodermal genes among DEGs at 2.5h, transcription factors *osr1, osr2, sox17a, sox17b* as well as *hhex* (Fig. 1H).

### Regulation of *tbx2, id3* and *prdm1* by GATA4

To validate identified DEGs as physiologically relevant targets of GATA4 we examined their regulation by GATA4 in vivo. As shown in Fig2B, downregulation of GATA4 using morpholino oligonucleotides (MOs) leads to upregulation of *id3* at gastrula and neurula stages; interestingly, even though *id2* and *id4* were not confirmed as DEGs, their expression was also upregulated by GATA4 MOs, suggesting an indirect mode of action. Downregulation of GATA4 and most closely related cardiogenic GATA factor, GATA5, led to upregulation of *prdm1*, and to downregulation of *tbx2* (Fig. 2E). This analysis has shown that GATA4 is required for normal expression of *id3, prdm1* and *tbx2* in vivo.

We next asked if GATA4 directly regulates *id3, prdm1* and *tbx2* through association with their regulatory regions. For this analysis we identified consensus GATA binding sites within 10 kb upstream and 2 kb downstream of their translation start sites and tested their presence in genomic DNA associated with exogenous GATA4. For all 3 genes, the minority of tested sites were found to associate with GATA4 (Fig. 2C, F and Fig S3B-C), suggesting that GATA4 is directly and specifically regulating their expression.

We subsequently tested if *tbx2* and *prdm1* have roles of their own in *Xenopus* heart development. The expression of *tbx2*, a GATA4 target upregulated by cardiogenic conditions, was downregulated by MOs, and this led to reduced and deformed heart formation in majority of embryos (Fig. 2G and Fig. S3D). The role of *prdm1* as a target of GATA4-mediated downregulation was tested by mRNA-mediated overexpression, which caused morphological defects such as cardia bifida and linear heart tube (Fig. 2G and Fig. S3E). These results indicate that *tbx2* and *prdm1* are targets GATA4 in cardiogenesis and that their normal expression is required for early heart development in *Xenopus* embryos.

### GATA4 is required for normal expression for *PRDM1* and *TBX2* during human iPSC-CM differentiation

We next sought to establish whether these findings are translatable to human development using iPSC-cardiomyocyte (iPSC-CM) differentiation to model human cardiomyogenesis. Firstly, the expression of *GATA4, PRDM1* and *TBX2* was examined during normal iPSC-CM differentiation. Consistent with previous studies *GATA4* expression is observed from the onset of differentiation with a noticeable increase in expression at day 4 (Fig. 3A), coincident with the establishment of the early cardiac progenitor (CP) pool (Churko et al., 2018). Its expression rises throughout differentiation peaking approximately when beating becomes apparent (day 9-10 of differentiation). Expression of *PRDM1* peaks early at day 3 and decreases when *GATA4* expression initially peaks at day 4 (Fig. 3A). Conversely *TBX2* expression becomes prominent only after the peak in *GATA4* expression. The expression patterns of these genes support the notion that the regulatory relationship between GATA4, *PRDM1*, and *TBX2* may be a conserved feature of cardiogenesis.

**Figure 3.**
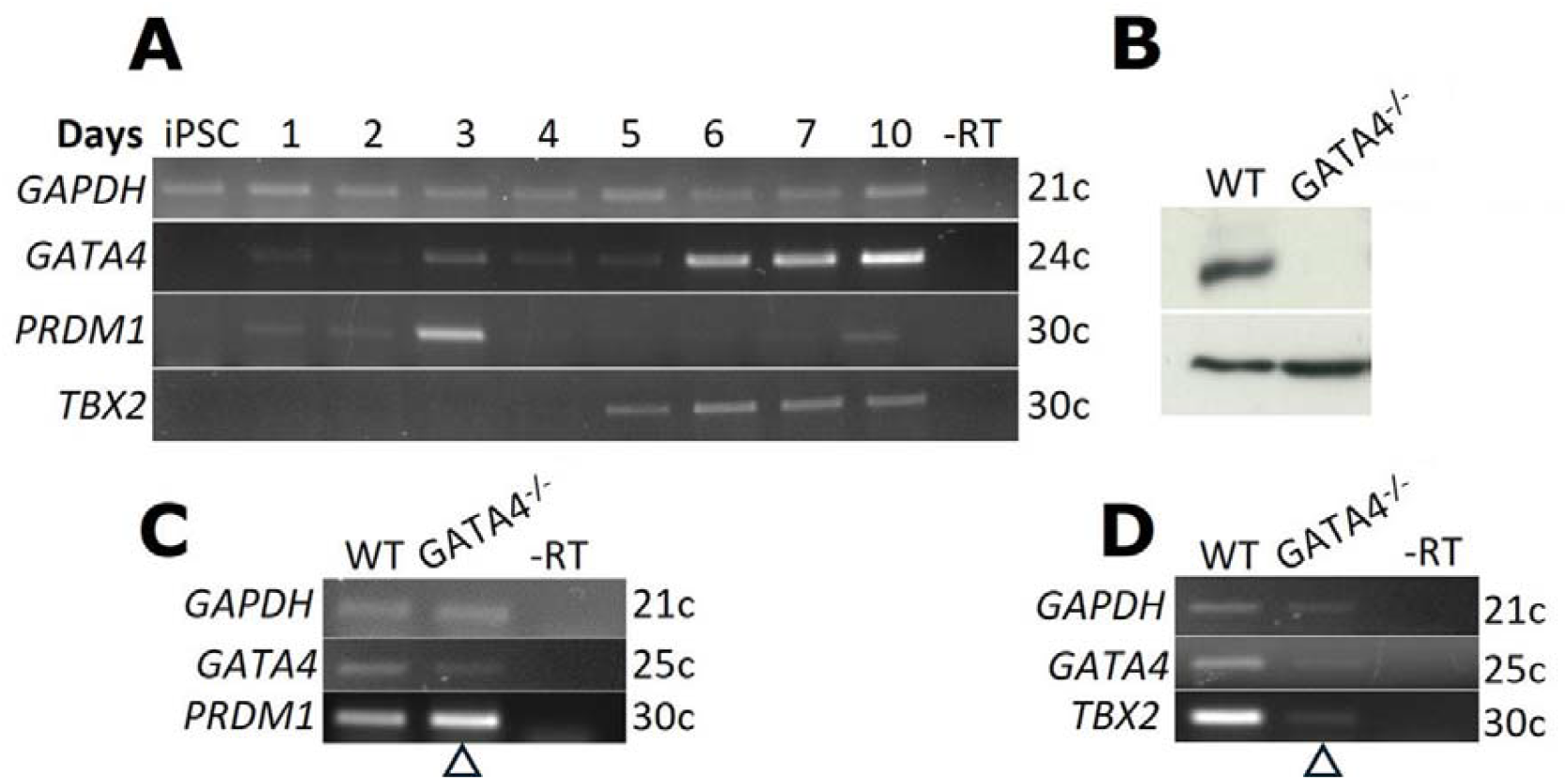
GATA4 is required for regulation of its target genes PRDM1 and TBX2 in iPSC-CMs. (A) Analysis of the expression of GATA4 and its target genes during iPSC-CM differentiation examined using RT-PCR. Representative of 3 repeats. (B) A western blot showing the absence of GATA4 protein at day 4 in the GATA4 null lines created using CRISPR-Cas9 gene editing. Representative of 3 repeats. ERK2 is used as a loading control. (C) RT-PCR data showing the expression of *GATA4* in mutant cells at day 4 is reduced, versus an increase in the expression of *PRDM1* (arrowhead), when compared to WT cells. (D) RT-PCR for day 10 samples showing *GATA4* expression remains lower in the mutant line, as is *TBX2* (arrowhead). Representative of 2 experiments. For all RT-PCRs *GAPDH* has been used as a normalisation control.

To establish whether the expression pattern of *PRDM1* and *TBX2* depends on GATA4, CRISPR-Cas9 gene editing was used to inactivate *GATA4* (Fig. S4). Knockout of the protein was confirmed by western blot (Fig.3B). RNA-sequencing and RT-PCR was then used to examine the expression of the target genes in the GATA4 null line. As shown in Fig. 3C-D, the expression of *PRDM1* and *TBX2* is disrupted in the absence of GATA4, with *PRDM1* expression being higher than normal, and *TBX2* expression being lower than normal. These results are consistent with the hypothesis that GATA4 regulates *TBX2* and *PRDM1* in cardiogenesis in humans as well as in *Xenopus*. Additional DEGs identified in the *Xenopus* screen showed dysregulation in the GATA4 null iPSC-CMs (Fig. 5H), suggesting that GATA4-regulated GRN has more extensive conservation besides TBX2 and PRDM1.

### GATA4 deficient iPS cells fail to form functional cardiomyocytes

The GATA4 null iPSC lines were differentiated using Wnt modulation protocols (Burridge et al., 2014; Lian et al., 2012) and were examined up to Day 32 of differentiation. No beating was observed in 12 independent differentiation experiments. Staining for TNNT2 by immunofluorescence was used to detect cardiomyocytes in iPSC-CM differentiation of WT and GATA4 null lines. On average there was a 4-8 fold decrease in the number of TNNT2 positive cells (TNNT2+) formed from GATA4 -/- iPSCs in comparison to WT (Fig. 4C). In the few mutant cells that expressed TNNT2 the signal also appeared weaker. Mean fluorescence per cell was used to quantify this and the average signal in the GATA4-/-TNNT2+ cells was found to be 2.5 times lower than for WT TNNT2+ cells (Fig. 4D). Examination of the cells at a higher magnification reveals that in the TNNT2+ GATA4-/-cells myofibrils lack a normal striation pattern (Fig. 4E, S6A). Consistent with the results seen for TNNT2, staining for additional sarcomeric markers MYBPC3 (Fig. 4E, S6B) and filamentous actin (Fig. S6C) was also disrupted in the GATA4 null lines. In contrast, staining for general cytoskeletal protein α-tubulin (TUBA1A) appears normal (Fig. S6D).

**Figure 4.**
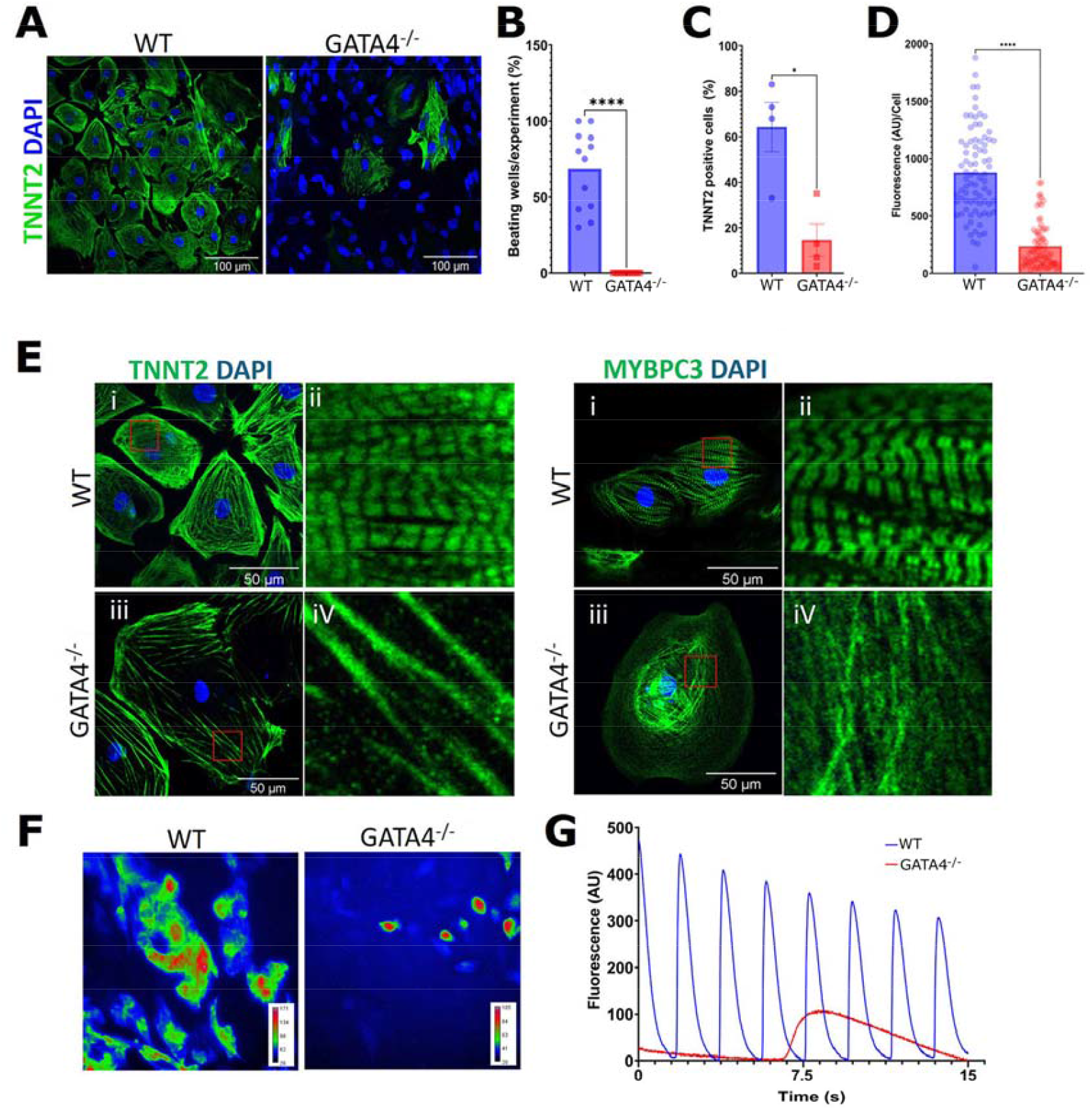
GATA4 null iPSCs fail to form normal cardiomyocytes. (A) Immunofluorescent (IF) staining for TNNT2 (green) in WT and GATA4^-/-^ mutant cells at Day 32 of cardiac directed differentiation. DAPI (blue) was used to counterstain nuclei. Representative examples from 4 experiments are shown. (B) A comparison of the proportion of beating wells detected per experiment between the WT and GATA4 null line. Based on 12 independent differentiation experiments. (C) A comparison of the proportion of TNNT2 positive cells identified by IF between the cell lines. TNNT2+ ive WT cells = 64.25% ± 10.87 and GATA4 -/- = 14.50% ± 7.14. (D) TNNT2 mean fluorescence per cell for WT and GATA4^-/-^, n = 3. For panels B and C a one-way ANOVA was used for statistical analysis. For panel D, a student’s t-test has been used. P-values: * ≤ 0.05, and **** ≤ 0.0001. (E) Representative images of individual WT and GATA4^-/-^ cells at 40x magnification stained for TNNT2 or MYBPC3 (green), DAPI (blue) was used to counterstain nuclei. The enlarged images outlined in red box provide a closer look at individual myofibrils. In the WT line >90% of the CMs present have clear sarcomeres compared to <7% for the GATA4-/-line. Further examples are presented in supplemental figure S6. (F) Fluctuations in calcium levels were measured over time in a field of WT and GATA4^-/-^ cells at Day 32 of differentiation using calcium indicator Fluo-8AM. The fluctuation in signal is shown as an RGB pseudo-coloured projection. Red areas being pixels with the highest signal variance over time, and blue being the lowest, n = 3. (G) In addition to these overviews, calcium flux was plotted over time from selected regions of interest for each cell line. For each cells line n= 9, with 3 wells analysed across 3 independent experiments.

The physiological activity of the WT and GATA4-/- cells was examined at Day 32 of differentiation using calcium imaging. Figure 4F shows that calcium flux can be observed across a large proportion of WT cells; conversely in the GATA4-/-cells this activity is restricted to a few small regions (Fig. 4F). When regions of interest for each are plotted individually it is apparent that calcium cycling in the WT population is frequent and regular (Fig. 4G). In the GATA4 null cells the transients detected were on average 6 times slower than their WT counterparts. This together with the low differentiation efficiency, rare and low TNNT2 expression, and lack of sarcomere formation in the cells lacking GATA4, accounts for the failure to form beating cardiomyocytes.

### GATA4 null cells likely adopt alternative cardiac cell fates

The vast reduction in the number of cardiomyocytes (TNNT2 and MYBPC3 positive cells) in the mutants raises the question of the identity of the remaining cell types.

To address how GATA4 null lines diverge from the WT path of differentiation, RNA-sequencing was carried out on cells at day 2, 5, and 10 of differentiation. At day 2 expression of GATA4 is low in the WT line therefore a large effect on the transcriptome at this time was not expected. Indeed, the GATA4 null cell transcriptome is comparable to that of the WT cells (Fig. S5A,B) as is the expression of early mesodermal and cardiac mesoderm markers *T* and *MESP1*, respectively (Fig. S5F).

At day 5 in the WT population the expression of cardiac progenitor genes including GATA4 rises. The levels of these genes are lower in the GATA4 null population and at day 10 the discrepancy between the WT and GATA4 null line becomes exaggerated. At day 10 the most affected cardiac GRN member is *TBX5*, being almost undetectable in the GATA4-/-cells (Fig. 5A,B). However, not all members of the cardiac GRN are downregulated: for example, *MEF2C* expression is expressed at a similar level in both lines and *ISL1* expression is increased, strongly suggesting that members of cardiac GRN display differential requirements for GATA4 function (Fig. 5A,B). Analysis performed at additional timepoints demonstrates that *TBX5* expression is undetectable during cardiomyocyte differentiation of the GATA4 null line (Fig. 5B. In contrast, expression of *NKX2-5* is readily detectable and expression of *ISL1* is higher after Day 6 (Fig. 5B).

During directed differentiation of iPS cells to cardiomyocytes other cardiovascular cell types are formed such as fibroblasts, endothelial, and smooth muscle cells (Grancharova et al., 2021; Jiang et al., 2022). Therefore, these cell types are the most likely candidates for the observed non-cardiomyocytes in GATA4 null mutant cell population.

**Figure 5.**
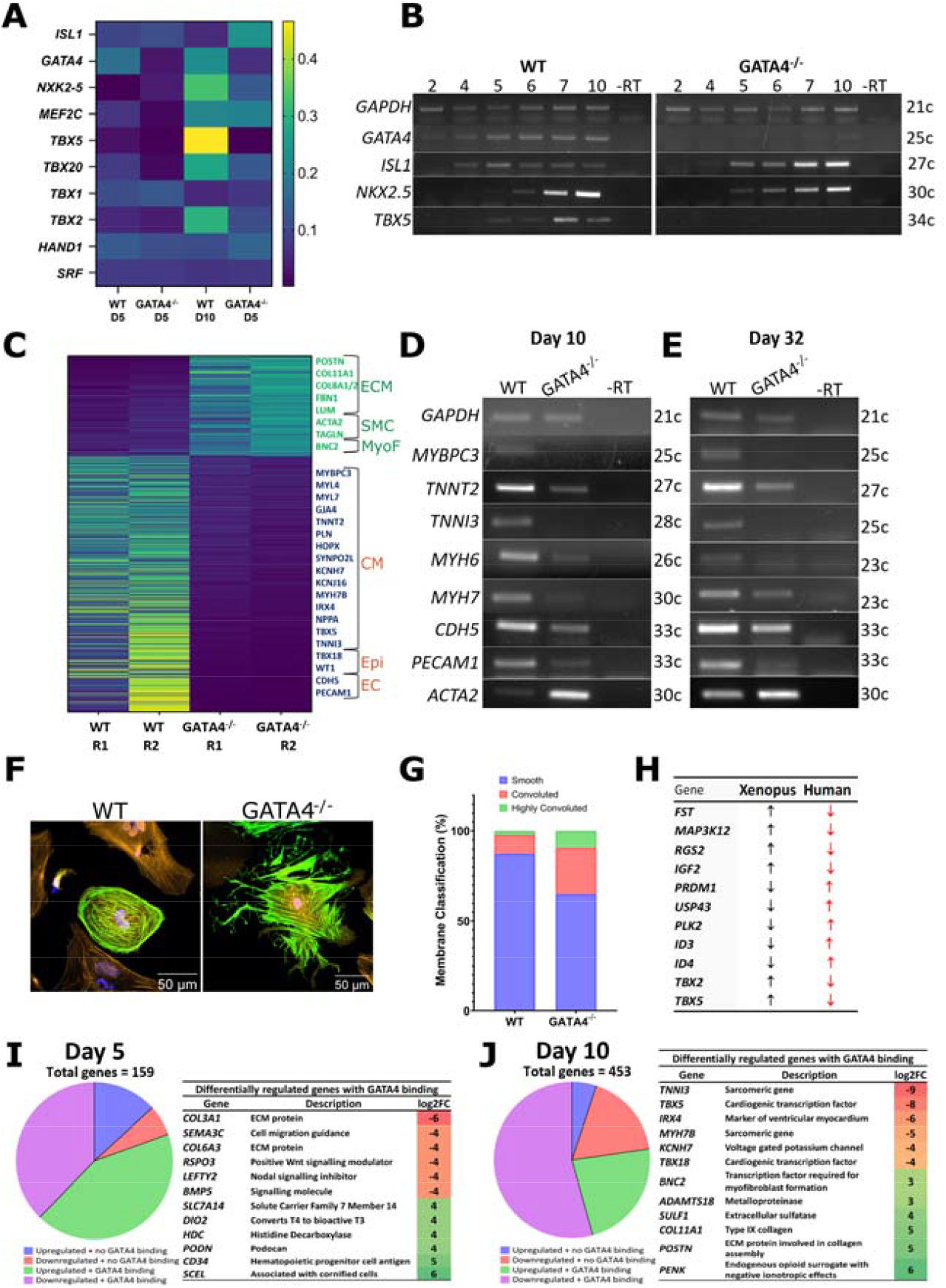
In GATA4 -/- cells, cardiomyocyte and endothelial cell transcriptomes are suppressed whereas smooth muscle cell and fibroblast genes are upregulated. (A) A heatmap showing the normalised expression of a selection of marker genes indicative of progression through iPSC-CM differentiation (B) Analysis of a number of cardiac progenitor marker genes by RT-PCR, expression of GAPDH was used for normalisation. n =2. (C) A heatmap showing the normalised expression of a selection of cardiomyocyte (CM), myofibroblast (MyoF), endocardial (EC), extracellular matrix (ECM) and smooth muscle cell (SMC) genes at Day 10. A selection of the genes from C were validated in an additional RNA sets at days 10 (D) and 32 (E) of differentiation using RT-PCR (n = 2). (F) Immunofluorescent images showing WT and GATA4 -/- cell at 40x magnification at day 32 of differentiation. Stained for TNNT2 (green), cell mask orange (Orange), and nuclei stained with DAPI (blue). (G) Cell membrane morphology was classified for WT and GATA4 -/- cells into smooth, convoluted, and highly convoluted. Those belonging to the convoluted category have membrane extensions, and those in the highly convoluted category displaying many of these branching extensions. WT cells analysed = 88, GATA4-/- = 55. (H) A table showing the direction of change for a selection of prospective GATA4 targets identified in the *Xenopus* screen (GATA4 overexpression) and their expression change in the GATA4 null iPSC line. Upregulated genes in the *Xenopus* column would be expected to be downregulated in human samples, and vice versa. (I-J) The pie charts displayed show the number of DEGs in the GATA4 mutants, at days 5 and 10 respectively, with a Log2FC value of 2 or more that are bound by GATA4 in foetal or adult mouse hearts, according to datasets generated by Akerberg *et al*. 2019. The accompanying tables list a selection of these bound genes and their log2FC value from our RNA-seq analysis of the GATA4-/- null line.

In agreement with previous report that has identified an essential requirement for GATA4 in the formation of the epicardium (Watt et al., 2004) a decrease in the expression of epicardial marker genes (*TBX18* and *WT1)* was observed (Fig. 5C). Furthermore, the expression of endothelial marker genes *CDH5* and *PECAM1* was decreased to a similar degree as what was seen for the cardiomyocyte specific gene set, suggesting that cardiomyocytes, epicardial cells and endothelial cells all require GATA4 for their development in this model (Fig. 5C-E). The expression of markers which are highly but not exclusively expressed in smooth muscle cells such as *VIM, TAGLN*, and *ACTA2* was upregulated in the GATA4-/- mutant cells (Fig. 5C-E). Marker of activated fibroblasts *POSTN* (Fig. 5C) was also upregulated, as well as numerous ECM components (Fig. 5C and Supplemental Table 2). Our gene enrichment analysis indicates significant over-representation of multiple ECM terms, consistent with upregulation of smooth muscle and fibroblast markers. Among the genes upregulated in GATA4 -/- cells at Day 10 is *BNC2*, a recently described master regulator of myofibroblast differentiation (Fig. 5C) (Bobowski⍰Gerard et al., 2022). Closer inspection of cellular morphologies of mutant TNNT2+ cells revealed an appearance of highly convoluted cells suggestive of a myofibroblast phenotype (Fig. 5F).

Together, these results suggest that the loss of GATA4 adversely affects the development of the cardiomyocytes and endothelial lineages, whilst simultaneously promoting the development of related mesenchymal cells-activated fibroblasts, myofibroblasts and smooth muscle cells.

To see if GATA4 could directly regulate DEGs, we analysed the high resolution GATA4 ChIP-seq data set for adult and foetal mouse heart (Akerberg et al., 2019). From our dataset at day 5, 159 DEGs were found to be consistently differentially regulated by a log2FC value ≥2. Of these genes 80% were associated with ≥1 GATA4 binding sites. At day 10 by these criteria 453 genes were found to be differentially regulated and of these 77% were found to be associated with ≥1 GATA4 binding sites (figure 5I-J). Importantly these included genes differentially expressed in both *Xenopus* and human RNA-seq datasets (Fig. 5H). Both up- and down-regulated DEGs were identified as direct targets based on GATA4 binding, in agreement with known activities of GATA4 in both transcriptional repression and activation.

### TBX2 is required for normal cardiomyocyte differentiation

We next examined whether the GATA4 target genes identified and validated in the *Xenopus* screen are themselves required for normal iPSC-CM differentiation. CRISPR-Cas9 gene editing of the TBX2 locus caused an in-frame deletion of DNA binding domain, likely resulting in a loss of function mutation (Fig. 6A,B, Fig. S8). The *TBX2* mutant iPS cell line produces cardiomyocytes at a similar rate to WT iPS cells (Fig. 6C). However, the cells produced are on average 1.6-fold larger than WT cells at day 12 of differentiation. This difference is accentuated at day 32, with the size of the mutant cells around 2.3-fold larger (Fig. 6D). Additionally, examination of the myofibril and sarcomeric structure of the *TBX2* mutant cardiomyocytes by immunofluorescence (Fig. 6F-H) reveals them to be significantly more disorganised (74%) than their WT counterparts (40%).

**Figure 6.**
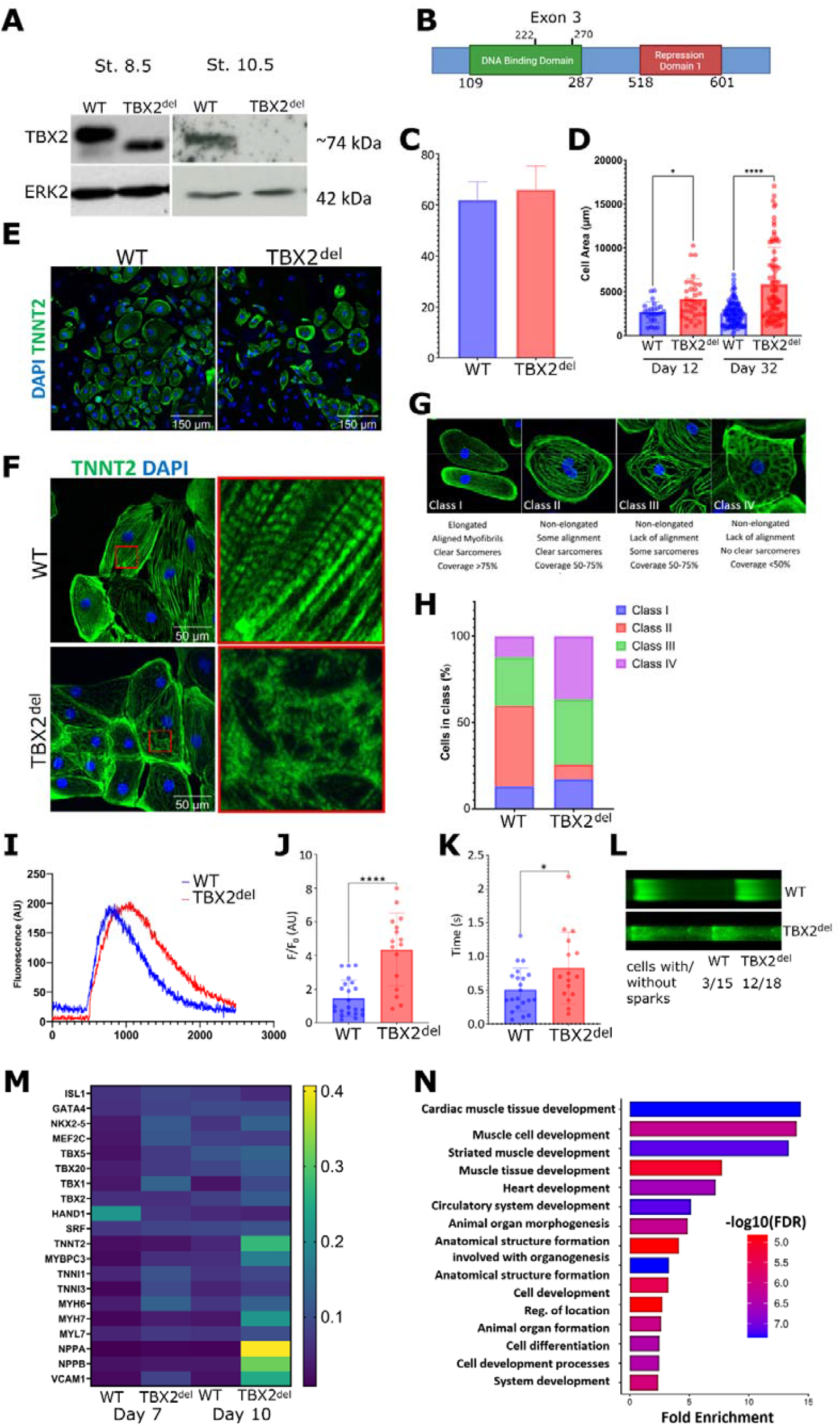
TBX2 mutant cells show pathological hypertrophy and upregulation of CM genes. To better visualise annotations in the figure, the full name TBX2^c.700_761del^ is abbreviated as TBX2^del^. (A) A western blot showing TBX2 expression in *Xenopus* embryos injected uniformly with WT TBX2 or TBX2^c.700_761del^ mutant mRNA at 2-4 cell stage. The embryos were harvested for protein analysis at stages 8.5 (blastula) and 10.5 (early gastrula). ERK2 was used as a loading control. The mutant protein is readily detected at st. 8.5 but is subsequently degraded at st. 10.5. (B) The TBX2 protein structure is shown with the DNA binding domain indicated in green. In TBX2^c.700_761del^ mutants, exon 3 is skipped. The region of the TBX2 DNA binding domain affected by this is indicated. Image produced using BioRender. (C) The proportion of TNNT2 positive cells in the total cell population was recorded. The values are given as a percentage ± SEM and are as follows; WT 62% ± 7.00 and TBX2^c.700_761del^ 66% ± 9.23 (n=3). (D) The surface area of the CMs was measured across 3 differentiations at day 32 and 1 differentiation at day 12. (E) Representative fields of WT and TBX2 mutant day 32 iPSC-CMs are shown at 40x magnification. The cells were stained for TNNT2 (green) to identify CMs, and counterstained with DAPI (blue) to identify nuclei. (F) Enlarged pictures of WT and TBX2 mutant CMs at day 32 showing disarray of myofibrils and sarcomeres. (G) Representative images of cells belonging to classes I-IV, with class I cells showing the highest level of order and class IV the most severe disorganisation. (H) Quantification of cardiomyocyte disorganisation in the WT and TBX2 mutant line through categorisation into one of the four categories described in (G), n =3. (I) Representative Ca^++^ transients for WT and TBX2^c.700_761del^ mutants at ∼ day 32 of differentiation for each cell line. (J) The amplitude of individual Ca^++^ cell transients. (K) Calcium imaging time to peak measurements. A One-way ANOVA has been used for all statistical comparisons shown. P-values; * ≤ 0.05, *** ≤ 0.001, and **** ≤ 0.0001. (L) (*upper panel*) Exemplar confocal Ca^++^ line scan recordings of Ca^++^ transients in a WT and TBX2 mutant cell. Spontaneous Ca^++^ sparks are visible during diastole between beats. (*lower panel*) TBX2 mutant cells tended to exhibit more diastolic Ca^++^ sparks than WT cells, although this was not statistically significant (P = 0.09; X^2^ test) (M) The heatmap displays the normalised RNA-seq data for a selection of CP and CM differentiation markers, n = 2. (N) A selection of GO terms found to be upregulated in the TBX2^c.700_761del^ mutant at day 10 of differentiation. Figure produced using ShinyGo.

In the *TBX2* mutants the amplitude of the calcium peaks recorded was notably higher, and there was a larger range of amplitudes observed indicating that there is a wider heterogeneity in the Ca^++^ handling properties of the mutant cardiomyocytes (Fig. 6 J-K). *TBX2* mutants also displayed increased propensity for formation of spontaneous calcium sparks, indicating defective calcium handling (Fig. 6L). Taken together these results indicate that interfering with TBX2 function promotes formation of maladaptive hypertrophic cardiomyocytes.

To further explore the phenotype of the cells, RNA-seq was conducted on samples taken at days 7 and 10. These time points correspond with the peak of *TBX2* expression during iPSC-CM differentiation and the onset of beating, respectively. Notable findings include increased expression at day 10 of *NPPA* and *NPPB* (Fig. 6M), genes whose elevated expression is known to be associated with heart failure (Tan et al., 2002). TBX2 is known to repress *NPPA* in AVC (Habets et al., 2002), thus increased *NPPA* expression in *TBX2* mutant iPSC-CMs strongly suggests that the mutation leads to loss of function. *VCAM1* is another gene that has been associated with heart failure and immune cell infiltration of the myocardium (Troncoso et al., 2021; Wang et al., 2021) and was also upregulated at day 10 (Fig. 6M). The increased expression of several sarcomeric genes e.g. *TNNT2, MYBPC3*, and *MYH7* in *TBX2* mutant cells is consistent with the hypertrophic phenotype (Fig. 6M). In agreement with this finding, GO terms related to cardiomyocyte development such as Cardiac Muscle Cell Development were enriched (Fig. 6N, Supplemental Table 2).

### PRDM1 deficient iPSC-CMs appear structurally unaltered but show premature beating

We next examined the requirement for *PRDM1* in iPSC-CM differentiation through the creation of a *PRDM1* null line using CRISPR-Cas9 gene editing (Fig. S10). Knockout of the protein was confirmed by western blotting (Fig. 7A). The *PRDM1*^*-/-*^ line produced CMs at a similar efficiency to WT and the phenotype of these cells when examined by IF resembles that of WT cells (Fig. 7 C-D). In the PRDM1-/-line beating started around 2 days earlier on average than in the WT line (Fig. 7B). Gene expression analysis of several sarcomeric genes such as *MYL7, MYH6, MYH7*, and *TNNI3* indicates subtle mis-regulation, early increase at day 7 and subsequent decrease relative to the WT cells (Fig. 7E). RNA-seq results show that at day 3 when the level of PRDM1 is highest, upregulated DEGs greatly outnumber downregulated DEGs (Fig. 7H) and include genes associated with neural development and pluripotency (Supplemental Table 2) suggesting that transcriptional repressor PRDM1 might be fine-tuning the timing and levels of gene expression during cardiomyocyte differentiation and repressing early alternative gene programs.

**Figure 7.**
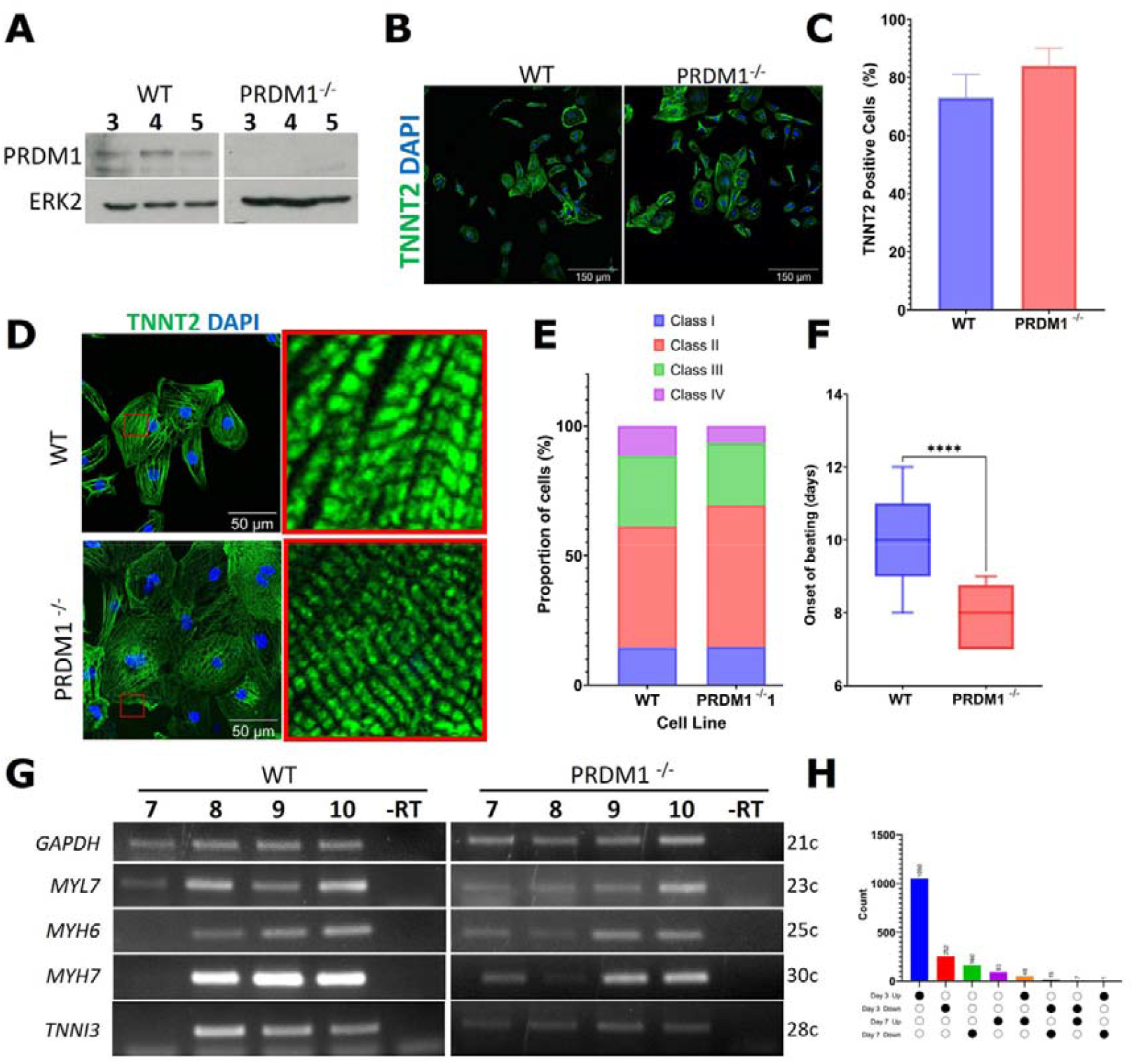
Accelerated formation of CMs from PRDM1^-/-^ iPS cells. (A) Western blot analysis of PRDM1 protein in WT and mutant iPS cells at indicated days of iPSC-CM differentiation. ERK2 is used as a loading control. (B) Beating onset was recorded for WT (average onset = day 9.9) and PRDM1^-/-^ (average onset = day 7.8) lines, n = 8. (C) CM differentiation efficiency, assessed by staining for TNNT2 (green) at day 10, is not affected in PRDM1^-/-^ cells. The values are given as a percentage ± SEM and are as follows; WT 73% ± 16.3 n=576; PRDM1^-/-^ 84% ± 12.3 n=367, across n = 3 experiments. (D) Representative images of WT and PRDM1^-/-^ cells stained for TNNT2 (green) and counterstained with DAPI (blue) for nuclei detection at day 32 of differentiation. Shown at 20x magnification. (E) Samples collected at day 7 of differentiation were subjected to RNA-seq. A heatmap showing the expression of cardiac progenitor and cardiomyocyte specific genes in the WT and PRDM1^-/-^ lines at day 7 of differentiation, n = 1. (F) Expression of a selection of sarcomere genes analysed by RT-PCR in a WT and PRDM1^-/-^ differentiation. *GAPDH* is used as a loading control. Days of iPSC-CM differentiation and number of PCR cycles are indicated. (G) An upset plot showing the number of DEGs per time point and their direction of change. (H) Comparable sarcomeric structure was seen between the WT and PRDM1 mutant cardiomyocytes. Staining for sarcomeric protein TNNT2 (green) in day 32 hCMs formed from WT and PRDM1 mutant cells, counterstained with DAPI (blue); enlarged images are shown on the right-hand side. (G) The proportion of cells in each class, WT n = 180 and PRDM1-/-, n = 150, over n = 3 experiments. The classification system utilised is as described in Figure 6.

## Discussion

Cardiac development relies on complex interactions of numerous transcription factors within the cardiac gene regulatory network. The knowledge of the composition of this network and how it operates is still incomplete. Here, using cardiogenic factors GATA4 and Nodal in pluripotent explants from *Xenopus* embryos we have examined the cardiogenic gene program. We identified a common gene set on which these cardiac inducing factors converge in order to regulate cardiogenesis. We focused on transcription factors *prdm1* and *tbx2* whose expression in X*enopus* embryos is regulated by GATA4, possibly directly. Using iPSC-CM differentiation and CRISPR-Cas9 gene editing we find this regulatory relationship to be conserved in human cardiomyogenesis and that target genes *PRDM1* and *TBX2* can modulate the cardiac gene program.

Our results demonstrate that GATA4 is essential for differentiation of human cardiomyocytes from iPS cells. Previous work in mouse, *Xenopus* and zebrafish models has established distinct stage-dependent roles for GATA4 during cardiomyocyte development, with defective heart still present in null mutant embryos (Borok et al., 2016; Crispino et al., 2001; Haworth et al., 2008; Holtzinger and Evans, 2005; Watt et al., 2004; Zeisberg et al., 2005). In the current study we show that human GATA4 -/- iPS cells lines have a greatly reduced capacity to form cardiomyocytes and do not show beating activity, a phenotype which is stronger than the phenotypes gained from GATA4 null mutation in animal models. Furthermore, a recent report by (Gonzalez⍰Teran et al., 2022) presented results for a human *GATA4* -/- iPS cell line that can form beating cardiomyocytes, albeit at a greatly reduced frequency. The reasons for this apparent discrepancy are not clear but likely include experimental details such as genetic background, which is known to be a major determinant of phenotypic variation in human iPSCs (Kilpinen et al., 2017).

The failure of GATA4 -/- cells to form beating cardiomyocytes in this study is underlined by a greatly reduced differentiation efficiency and defective sarcomere formation. Genome wide transcriptomic analysis has documented the downregulation of cardiomyocyte and endothelial cell gene sets, and an upregulation of genes associated with smooth muscle and activated fibroblast cell phenotype. These include a marker of activated fibroblasts *POSTN* (Shimazaki et al., 2008) and recently described master regulator of myofibroblasts *BNC2* (Bobowski⍰Gerard et al., 2022). A subpopulation of GATA4 null cells displayed myofibroblast-like convoluted shape but was also found to be positive for TNNT2. Expression of cardiomyocyte-specific sarcomeric genes *TNNT2* and *MYBPC3* has been identified in a subpopulation of human cardiac fibroblasts (Litviňuková et al., 2020). It will be of interest to investigate if loss of GATA4 promotes transition of those fibroblasts to myofibroblast phenotype.

Loss of GATA4 also upregulates numerous genes involved in fibrosis and inflammation such as *COL3A1, COL8A1*, and *COL11A1, IL11* as well as *IGFBP7*, which is associated with senescence and heart failure (López et al., 2021; Moschen et al., 2011; O’Reilly, 2023; Xiang et al., 2017; Zhang et al., 2022).

The impact of GATA4 deficiency on cardiac GRN was not uniform: whilst the expression of *TBX5* was undetectable, the expression of *ISL1* (at Day 10) and *NKX2-5* (at Day 5) was upregulated in GATA4 -/- cells. This finding is consistent with the proposed repression of *NKX2-5* by GATA4/5/6 in vivo (Jiang et al., 1999). Of note, *NKX2-5* is known to be associated with ECM remodelling and pathological smooth muscle cell phenotype (Papaioannou et al., 2024). It will be of interest to determine if *NKX2-5* has a similar role in regulating pro-fibrotic phenotype in GATA4 -/- iPSC-CM cells.

The downregulation of *TBX18* and *WT1* observed in GATA4 -/- iPSC-CM cells suggests defective epicardial development, in agreement with an in vivo requirement of GATA4 for development of the epicardium in the mouse embryo (Borok et al., 2016; Watt et al., 2004). This is an apparently paradoxical observation as select markers of epicardial derivatives, smooth muscle and fibroblasts, are upregulated in GATA4 mutant human iPSC-derived cells. One possibility is that diminished epicardial development still permits formation of abnormal smooth muscle cells and fibroblasts.

GATA4 is known to be expressed in cardiac fibroblasts. Our results show that deletion of GATA4 leads to pro-fibrotic and myofibroblast gene expression signature, suggesting that it is required for the normal fibroblast phenotype. In agreement with this hypothesis, overexpression of GATA4 in cardiac fibroblasts was recently shown to reduce heart failure-induced fibrosis (Yamada et al., 2024). The role of GATA4 in attenuating fibrosis is not restricted to cardiac fibroblasts, as it also induces regression of liver fibrosis by deactivating hepatic stellate cells (Arroyo et al., 2021).

A potential mechanistic basis for the observed features of smooth muscle and activated fibroblasts in GATA4 null cells is offered by a recent study by Robbe *et al*. (Robbe et al., 2022) which demonstrated that GATA4 physically interacts with repressive chromatin modifier CHD4 recruiting it to non-cardiomyocyte gene regulatory sites in mouse myocardium. The deletion of a GATA site in the modulatory region of smooth muscle gene Myh11, led to its upregulation in cardiomyocytes (Robbe et al., 2022). Misexpression of Myh11 in cardiomyocytes has previously been shown to result in sarcomere disarray, and results in a significant decrease in ventricular output (Wilczewski et al., 2018). Thus, it is possible that removal of GATA4 leads to de-repression of non-CM genes which based on our RNA-seq data are part of gene expression signatures of smooth muscle cells and activated fibroblasts. Future work will determine the precise nature of non-cardiomyocytes which express these genes.

Our results show that besides being a conserved target of GATA4 during cardiogenesis, *TBX2* is required for normal development of human iPSC-CMs. Mutant *TBX2* iPSC-CMs show hallmarks of pathological hypertrophy at the cellular and molecular levels. In addition, whole genome transcriptome analysis shows upregulation of cardiomyocyte stress genes as well as of cardiomyocyte differentiation gene sets. These features are consistent with the described role of TBX2 as transcriptional repressor of the myocardial gene program: in the absence of WT TBX2 activity, tight regulation of cardiomyocyte differentiation might be disrupted to result in upregulation of cardiomyocyte gene expression, leading to hypertrophy and stress phenotype.

The actions of TBX2 in cardiac development have been well characterised in the context of AVC and OFT formation, where it represses the myocardial gene program (Aanhaanen et al., 2011; Christoffels et al., 2004; Harrelson et al., 2004; Singh et al., 2012). However, *TBX2* is broadly expressed in the linear heart tube before becoming restricted to these specialised tissues, indicating a potential for a broader role in early cardiac development outside of the OFT and AVC. In mouse *TBX2* KO models AVC and OFT development is defective, whilst there was no effect on the development of myocardium (Harrelson et al., 2004; Singh et al., 2012). It is possible that the apparent discrepancy is caused by species-specific difference or by the difference between in vivo and in vitro models. The Wnt modulation method of cardiac differentiation used in this study primarily produces ventricular-like cardiomyocytes, with some atrial and node-like cells also present (Lian et al. 2013; Burridge et al. 2014b; Galdos et al. 2022). With differentiation protocols for the derivation of additional cardiomyocytes subtypes such as atrial, node-like, or AVC and conduction-like (Cyganek et al., 2018; Lyra⍰Leite et al., 2022; Prodan et al., 2022; Wiesinger et al., 2022; Ye et al., 2024) it will be possible to more precisely dissect any cardiac cell type-specific roles of human TBX2.

Loss of *PRDM1* expression does not have a major impact on the outcome of cardiac differentiation in human iPSC-CMs, as efficiency of differentiation and cardiomyocyte morphology are unchanged; instead, it is characterised by subtle differences in beating onset. In the absence of *PRDM1*, beating is accelerated by on average 2 days. In mutant cells, most differentially expressed genes are upregulated at Day 3 of differentiation and include genes associated with neural development. There are multiple examples of PRDM1 repressing differentiation in other developmental contexts, for example in mammary luminal and sebaceous gland stem cells, in migrating primordial germ cells, and enterocyte maturation (Ahmed et al., 2016; Harper et al., 2011; Horsley et al., 2006; Yamashiro et al., 2016). Therefore, in this scenario it is possible that the absence of PRDM1 leads to earlier cardiomyocyte beating through a lack of repressive activity, which might be required for tightly regulated progression through iPSC-CM differentiation.

In conclusion, we have established a conserved cardiac regulatory module of GATA4 and its target genes *PRDM1* and *TBX2*, identified in *Xenopus*, which also operates in the development of human cardiomyocytes.

## Limitations

A clear limitation of the current study is the lack of high-resolution information on cell fate trajectories in GATA4 -/- iPSC-CM cultures, which will be essential for better insight into the roles of GATA4 in specification and differentiation of cardiovascular cell types.

## Materials and Methods

### *Xenopus* embryos and explants

All work with *Xenopus laevis* was approved by Cardiff University’s Animal Welfare and Ethical Review Board and was undertaken under a license from the UK Home Office. *Xenopus laevis* embryos were obtained by mating of frogs primed with human chorionic gonadotrophin (Sigma) or by in vitro fertilisation (Sive, 2000). Jelly membrane was removed with 2% cysteine-HCl, pH 7.8 (Sive, 2000). Embryos were grown in 10% Normal Amphibian Media (NAM) and staged as described (Sive, 2000). Whole embryos (WE) or explants were cultured until age match control siblings had reached desired stage. Microinjections were carried using an IM 300 Micro-injector (Narishige Scientific), in 75% NAM containing 3% Ficoll (Sigma). Morpholino Oligonucleotides (MOs) were supplied from Gene Tools (http://www.gene-tools.com/) and injected at 10 nL/embryo. Tbx2 MO was described (Cho et al., 2011). GATA4 and GATA5 translation- and -splicing blocking MOs were described (Haworth et al., 2008). mMESSAGE mMACHINE kit (Ambion) was used for capped mRNA synthesis. Previously described templates for making capped mRNA: GATA4-GR (Afouda et al., 2005; Latinkić et al., 2003), Gata4 (Gallagher et al., 2012), *Xenopus* n*odal5* (Takahashi et al., 2000). *Xenopus prdm1* (de Souza et al., 1999) was obtained from the European *Xenopus* Resource Centre (Portsmouth, UK), and coding sequence was amplified by PCR and cloned into BamHI-XbaI sites of pCS2+ by PCR. WT and mutant human TBX2 CDS was cloned into pCS2+ using PCR. Correct sequence was confirmed by sequencing. Injection solutions included lineage tracers biotin- and rhodamine-dextran (Invitrogen; (Latinkić et al., 2003)). For cardiogenic Nodal signalling we injected 50-110 pg of *nodal5* mRNA and for lower, non-cardiogenic Nodal signalling we used 16 units of activin protein provided as XTC cell line conditioned media (Smith et al., 1988). Cardiogenic nodal5 and non-cardiogenic but muscle-inducing activin doses were confirmed (Fig. S1A).

### Chromatin Immunoprecipitation

Chromatin Immunoprecipitation was performed as described by Blythe et al. (Blythe et al., 2009) with minor changes: 2-300 embryos were injected at 2-cell stage with 0.4 ng of capped mRNA coding for HA tagged rat GATA4 (Gallagher et al., 2012). Stage 9-9.5 embryos were fixed in 50 mL falcon tube for 30min in 25 mL of 10% NAM, 1% formaldehyde on ice without agitation, briefly washed in 10% NAM and quenched for 10 min. in 25 mL 10% NAM, 0.125 M glycine followed by 3 washes in 10% NAM. Embryos were transferred into 1.5 mL microcentrifuge tubes (50/tube) and frozen on dry ice before storage at -80 ^0^C.

The crosslinked 50 embryos were thawed on ice for 15 min and homogenised in 600 mL of RIPA buff + Protease Inhibitor tablets (complete EDTA-free, Roche) and incubated for 15 minutes beginning from the time of initial homogenization. Homogenates were centrifuged for 5 min 3000 rpm, 4^0^C. Supernatant was removed and the inside wall of the tube above the pellet wiped with fine paper to remove lipid residue. The pellet was resuspended in 400 mL of RIPA buff +PI to get ∼450 mL. Sonication was performed with Soniprep 150 at 70% power: 8 cycles (10 sec. on – 50 sec. off) in 2 mL round bottom Eppendorf tubes on ice water bath. Shear median DNA size is ∼500 bp. The volume was adjusted to 650 mL with RIPA+PI and samples centrifuged for 10 min at 13000g, 4^0^C. Supernatant: 600 mL taken for IP, 5 mL taken for the INPUT, 20 mL for shear control. Anti-HA antibody magnetic beads were used according to manufacturer recommendations (Pierce). 50 µL of beads per 50 embryos, prewashed 3x 5min in TBS + 0.01% BSA using a magnet stand. Supernatant 600 mL taken for IP mixed with prewashed beads and incubated overnight rotating at 4^0^C. Supernatants were removed and beads stored at -20^0^C. Beads were washed 3x for 5 min in TBS-Tw (0.01% Tw20). Reversing crosslinks: beads (with chromatin), input and shear control samples were supplemented with 200 mL of TE buffer + 1/20 vol. of 5M NaCl and incubated overnight at 65^0^C. 1 mL RNAse A 10mg/mL was added to each sample and incubated for 30 min at 37^0^ C. + 1 mL Proteinase K (Roche), incubation 1h at 55^0^C. Proteins were removed by phenol-chloroform extraction and DNA precipitated with isopropanol. Samples were dissolved in 50 mL of ultrapure distilled water. PCR reactions were performed as for RT-PCR (below). Amplicons were chosen within 10kb upstream and 2kb downstream of translation start site based on presence of at least one consensus GATA binding site

### RNA extraction and RT-PCR analysis

RNA was extracted using the acid-guanidinium thiocyanate phenol chloroform method (Chomczynski and Sacchi, 1987) and cDNA was synthesised using the RevertAid reverse transcriptase kit (ThermoFisher). All primer sequences used were designed using NCBI’s primer blast program (Ye et al., 2012) and are listed in Supplemental Table 4. Samples were normalised using *odc1* primers for *Xenopus* samples and *GAPDH* primers for human samples. To find the appropriate amount of cDNA to use in further PCR reactions. PCR reactions were conducted in a volume of 25 μL, to each was added 0.1 μM of forward and reverse primer (Eurofins), 1X MyTaq red reaction buffer (Bioline), 0.5U KAPA Taq Polymerase (both KAPA Biosytems), and the appropriate volume of cDNA. An MJ Mini Thermal Cycler Machine (Biorad) was used for cycling. The amplicons generated were visualized using agarose gel electrophoresis. All primer sequences are presented in Supplemental Table 4.

Whole mount in situ hybridisation to detect expression of *myl7* in *Xenopus* embryos was performed as described (Gallagher et al., 2012).

### Human iPS cell culture and gene editing

All cell lines produced in this study were derived from Rebl Pat iPSCs kindly provided by Prof. Chris Denning, Nottingham University, and described in (Hammad et al., 2016).

### iPSC culture

iPSCs were cultured on Corning™ cell culture plates coated with Geltrex™ (ThermoFisher) at 30-45 µg/mL in DMEM/F12 Media (ThermoFisher) and maintained with B8 media formulated as described by (Kuo et al., 2020) . Passaging was carried out when cultures reached 75-80% confluency. ReLeSR™ was used to dissociate iPSC colonies as per the manufacturer’s instructions (StemCell Technologies), following this the cells were replated and fed with B8 2.5 μM Y27632 (HelloBio) for 24 hours following passaging before being switched to B8 for regular feeding every other day. All cells were maintained at 37°C in a humidified atmosphere with 5% CO_2_.

### CRISPR-Cas9 gene editing of iPSCs

All crRNAs were designed as pairs using CCTop (Stemmer et al., 2015). Each pair was intended to be used together to induce a frameshifting mutation. To produce gRNAs each crRNA (Integrated DNA Technologies) was combined with tracrRNA (Integrated DNA Technologies) at an equimolar concentration in IDTE Nuclease free buffer (30 mM HEPES (pH 7.5), 100 mM potassium acetate), then heated to 95°C for 2 minutes followed by gradual cooling to RT. Each gRNA was then combined with an equal volume of 6.2 μg/μL Alt-R S.p. HiFi Cas9 Nuclease V3 diluted in Cas9 storage buffer (10 mM Tris-HCl (pH 7.4), 300 mM NaCl, 0.1 mM EDTA, 1 mM DTT) and incubated together for 10 minutes at RT to form a ribonucleoprotein complex (RNP). Transfection of the RNPs into iPSCs was carried out in suspension using the P3 Primary Cell 4D-Nucleofector™ X Kit L (Lonza) and a Lonza 4D Nucleofector with program CA137. Single clones were selected and screened using gDNA PCR followed by gel electrophoresis. All mutations were confirmed using Sanger sequencing (Eurofins) and SyntheGo ICE (Conant et al., 2022) signal deconvolution. Sequences of crRNAs and genotyping PCR primers are provided in Supplemental Table 4. Following gene editing the pluripotency of the lines was assessed by confirming expression of *NANOG, POU5F1* and *SOX2* (Fig. S4, S5, S7, and S10).

### Karyotyping

Genomic DNA from the parental Rebl Pat human iPSC line and gene edited lines derived from it was extracted from >2 million cells using the Blood and Tissue DNA Extraction Kit (Qiagen), following the manufacturers guidance. The gDNA was karyotyped using the Infinium Global Screening Array-24, covering 654,027 markers across the genome at the MRC Centre for Neuropsychiatric Genetics and Genomics, Cardiff University. The results are summarised in Supplemental Table 5.

### CM differentiation and maintenance

Differentiation of iPSCs into CMs was carried out using two commonly utilised Wnt modulation protocols (Burridge et al., 2014; Lian et al., 2012). For differentiation cells were seeded at a density of 14 x 10^4^ – 17.5 x 10^4^ cells per cm^2^, in wells coated with Geltrex™ as above and expanded to 75-80% confluency in B8 medium before differentiation was initiated. For the CDM3 protocol published by Burridge *et al*. the cells were changed to CDM3 (RPMI 1640, 500 µg/mL, 213 µg/mL L-Ascorbic acid-2-phophate, Penicillin – Streptomycin 50 U/mL) with 6 µM CHIR99021 (HelloBio) for 48 hours. Following this media was changed to CDM3 with Wnt 2 µM WntC59 (HelloBio) for 48 hours. After these treatment stages cells were maintained in CDM3 media and refreshed every other day. Then switched to CDM3-L (RPMI 1640 no glucose, 500 µg/ml Fraction V BSA, 213 µg/ml L-Ascorbic acid-2-phosphate, 5 mM Sodium DL-Lactate) from days 10-14 to metabolically enrich the cultures for CMs.

For the GiWi protocol by Lian *et al*. the cells were plated as above and changed to RPMI B27^-insulin^ (ThermoFisher) with 6 µM CHIR99021 for 24 hours to initiate differentiation. This media was then removed and replaced with RPMI B27^-insulin^ for 48 hours. Following this the spent media was removed and mixed 1:1 with fresh RPMI B27^-insulin^ to make ‘conditioned media’.

To the conditioned media was added 2 µM WntC59 and the cells were maintained in this for a further 48 hours. After this the media was refreshed every 2 days with RMPI B27^-insulin^, and then with RPMI B27 from day 8-10. For cells differentiated using the GiWi protocol RPMI B27-L (RPMI 1640 no glucose, 1x B27, 5 mM Sodium D-L-Lactate) was used to metabolically select cells from day 10-14.

Upon finishing selection CMs derived using the CDM3 or GiWi protocols were maintained in CDM3 with media changes every 2 days.

### CM dissociation

The cell layer was washed with Hanks Buffered Saline Solution without Ca and Mg (HBSS, Merck). This was then removed and pre-warmed dissociation solution (HBSS, 200 U/mL Collagen Type II (Worthington Biotech), 1 mM HEPES pH 7.4, 10 µM Y37632 (HelloBio) was added and the cells incubated at 37 °C for 3-3.5 hours. The cell solution was then gently triturated combined with an equal volume of media and then filtered through a 100 µm filter. The cells in suspension were spun down at 400 rcf for 5 minutes, and the supernatant removed. The cell pellets were resuspended at an appropriate concentration in CDM3 with 10% heat inactivated FBS for re-plating. The following day the media was changed to CDM3.

### Immunofluorescence

Any cells to be analysed were dissociated as described above and then plated onto 13 mm glass coverslips coated with 30-45 µg/mL Geltrex™. The cells were allowed to attach and recover for a minimum of 48 hours before any further processing. The fixation and permeabilization method used was dependent on the antigen being detected (see Supplemental Table 3): either being carried out using 4% PFA for 10 minutes at RT, followed by permeabilization with 0.25% Triton-X-100 DPBS ^-Ca -Mg^ for 10 minutes at RT, or using 100% ice cold methanol for 15 minutes at 4⁰C. Regardless of fixation and permeabilization method used blocking was carried out with DPBS ^-Ca -Mg^ 10% serum for 1-hour at RT, with serum type matched to the secondary antibody used. Primary antibody incubation was carried out overnight at 4°C in blocking buffer, see Supplemental Table 3 for dilutions. The samples were then washed 5 times for 15-minutes at RT with PBS-0.1% Triton-X100, then incubated with 1:1000 Goat Anti-Mouse or anti-rabbit Alexa fluor conjugated antibodies (ThermoFisher) in blocking buffer overnight.

Hoechst 33342 at 1 µg/mL was used to counterstain nuclei by incubation for 15 minutes, followed by 3 five-minute washes in PBS ^-Ca2+ Mg2+.^ This was followed by mounting in Vectashield hard set mounting media with Phalloidin TRITC. Alternatively, HCS Cell Mask Orange (Invitrogen) was used at 1x to stain cell membranes and nuclei for 30 minutes, followed by three five-minute washes PBS ^-Ca -Mg^. Mounting was then carried out using Vectashield with DAPI (Vector Laboratories) to stain nuclei. A Zeiss LSM880 Confocal Microscope was used for imaging and FIJI/Image J Software (Schneider et al., 2012) for image analysis and editing. Following IF staining and imaging the TNNT2+ cells were analysed against criteria adapted from (Ang et al., 2016).

### Calcium Imaging

Any cells to be analysed were dissociated as described above and then plated onto 13 mm glass coverslips coated with 30-45 µg/mL Geltrex™. The cells were allowed to attach and recover for a minimum of 48 hours before any further processing. For analysis of the TBX2 lines and their WT counterparts’ cells were loaded with the Ca^2+^-sensitive indicator Fluo-4-AM (5 µmol/L) for 30 min at room temperature, followed by at least 10 min for de-esterification. Experiments were performed in a modified Tyrode’s solution containing, in mM: 133 NaCl, 5 KCl, 1 NaH_2_PO_4_, 10 4-(2-hydroxyethyl) -1piperazineethanesulfonic acid (HEPES), 10 glucose, 1.8 CaCl_2_, 1 MgCl_2_), pH 7.4 with NaOH. Confocal line-scanning microscopy was performed using an inverted Leica SP5 confocal microscope with a 63x 1.2 NA water immersion objective. Spontaneous Ca^++^ transients were recorded using the 488 nm line of an argon laser in line-scanning mode at 400 lines per second, with fluorescence emission collected between 500-650 nm. The confocal pinhole set to <2 Airy units. For the GATA4 lines and WT controls imaging was conducted using the widefield setting on the Olympus IX71 microscope. Before imaging the cells were loaded with 1 µM Fluo8-AM or 1 µM Fluo4-AM in cell culture media for 30 minutes at 37 °C. This media was then removed and Tyrode’s solution with 1mM HEPES pH 7.4 (Merck, T2397) was added. The cells were then imaged using a 488 nm laser. For both, temperature was maintained at 37 °C during Ca^++^ imaging experiments using a heated microscope enclosure. The change in fluorescence over time was plotted using FIJI/Image J Software (Schneider et al., 2012), and Ca^++^ transient analysis was performed using custom written MATLAB scripts. As part of this analysis non-cell background fluorescence was subtracted from any readings, then average cell fluorescence (F) was divided by the diastolic fluorescence (F_0_) to account for possible variability in dye loading. This was used to calculate an average for Ca^++^ transient amplitude value (reported as F/F_0_) and full duration at half maximum (FDHM).

### RNA-seq

The *Xenopus* RNA samples were converted to cDNA libraries using a Bioo NextFlex directional v.2 kit and were sequenced at Bristol University’s sequencing facility. The human RNA samples were converted to cDNA libraries using Illumina Truseq stranded mRNA library kit and sequenced at the Genome Research Hub, School of Biosciences, Cardiff University.

### RNA-seq Quantification and Differential expression

*Xenopus laevis* sequence analysis: paired end reads were aligned to *X. laevis* gene models v7.2 by JGI with Bowtie (Langmead et al., 2009) allowing for up to 3 mismatches in seed and leaving the other parameters at the default value. Gene hits were counted using RSEM (Li and Dewey, 2011)with default parameters. Expected read counts reported by RSEM were used to call differentially expressed genes which were called using DESeq (Anders and Huber, 2010). The dispersion was estimated with “per-condition” method and “gene-est-only” sharing mode. Each treatment was separately compared to control at each timepoint, genes with padj < 0.1 were defined as differentially expressed. To determine a gene’s differential expression irrespective of timepoint, we took the timepoint with the minimum padj.

Human sequence analysis: RNA-seq data (75bp single-end) data generated from Human iPSC cell lines (40-45 M per sample) were quality trimmed using fastp (Chen, 2023) and trimmed sequences aligned against the annotated Ensembl GRCh38.p14 human genome using STAR (Dobin et al., 2013). Quantification against gene objects was performed by using Rsubread (Liao et al., 2019) and differentially expressed genes (DEGs) derived using DESeq2 using the SARTools wrapper using default parameters (Varet et al., 2016). As with *Xenopus* data, each time point was compared to time and batch-matched controls and genes with p-adj <0.1 were defined as differentially expressed.

### Gene set enrichment

*Xenopus laevis:* to assess the composition of each group of genes we performed gene set enrichment using Enrichr. We took genes from each cluster with a known *Xenopus laevis* gene symbol and converted these to human symbols, by removing any “.N,” “-a,” or “-b” suffix for an integer N and converting to uppercase. We then made the following substitutions to convert certain known *Xenopus* gene symbols to human where the name of the ortholog has diverged or only a paralog exists: pou5f3 → POU5F1, mix1 → mixl1, dppa2 → DPPA4, lefty → lefty2, ventx1-3 → NANOG, mespb → MESP1, sox17a/b → SOX17. We remove any duplicate names that arose in this process. Gene nomenclature for *Xenopus laevis* we used is described at xenbase.org (Fisher et al., 2023) . For simplicity, in figures we omitted the information on homeologues (.L or .S) with the exception of *lefty1*.*S*, which was validated as DEG, unlike *lefty1*.*L*. We calculated enrichments for the following gene sets: KEGG_2021_Human, WikiPathway_2023_Human, BioPlanet_2019, GO_Biological_Process_2023, GO_Molecular_Function_2023, GO_Cellular_Component_2023. All enrichments are found in Supplemental Table 1.

Human Gene Ontology enrichment analysis was carried out using ShinyGO v0.77 and STRING v11.5 for network analysis (Ge et al., 2020; Szklarczyk et al., 2019) and is presented in Supplemental Table 2. Normalised counts for DEGs identified in human iPSC samples are presented in Supplemental Table 3.

### Western blotting

Cells were lysed in RIPA Buffer (50 mM Tris-HCl pH8, 150 mM NaCl, 0.1 % SDS, 0.5% Na Deoxycholate, 1 % NP-40, and 1x Protease Inhibitor Complete Mini (Roche)). Total protein extracts were mixed with 2x Laemmli Loading Buffer (BioRad) and boiled for 10 minutes before loading, 20 µg of total protein was used per well. Bands were resolved via polyacrylamide gel electrophoresis; the samples were then transferred to polyvinylidene difluoride membrane (Millipore). 5% Skimmed Milk in TBS Tween 20 (TBS-Tw: 5 mM Tris-Base pH 7.4, 20 mM NaCl, 0.1% Tween 20) was used to block the membranes for 1 hour. All antibodies used are described in Supplemental Table 4. For detection of TBX2 skimmed milk concentration was reduced to 1 % to improve signal intensity. Incubation with primary antibodies was carried out overnight in blocking buffer at 4°C with constant rotation. Following this the membranes were washed with TBS-Tw 3 times for 10 minutes each time. Secondary antibodies were then applied to the membranes diluted in 5% skimmed milk TBS-Tw or for detection of TBX2 1% skimmed milk TBS-Tw. The membranes were 3 times for 10 minutes each time in TBS-Tw to remove excess antibody. Clarity Western ECL (Biorad) was used for development followed by exposure of the membranes to Amersham Hyperfilm (Merck).

## Data Availability

The sequence data will be made publicly available in NCBI’s Gene Expression Omnibus through GEO Series accession numbers GSE308990 (human) and GSE308764 (*Xenopus*) as of the date of publication.

## Acknowledgements

This work was supported by grants from British Heart Foundation (PG/11/115/29287 and FS/18/42/33827) and from Heart Research Wales to BL. EDF is supported by a British Heart Foundation Intermediate Basic Science Research Fellowship (FS/IBSRF/21/25071). NO is supported by a Wellcome Career Development Award (227357/Z/23/Z). We are grateful to Prof. Chris Denning and his group (Nottingham University) for generously providing training in iPSC-CM techniques and for providing iPSC lines. We thank Prof. Nick Allen and his group for local support for iPSC culture work. Expert technical support by Alexandra Evans (MRC Centre for Neuropsychiatric Genetics and Genomics, Cardiff University) and Angela Marchbank (Genome Research Hub, School of Biosciences, Cardiff University) is gratefully acknowledged.

