## Supplemental figures for "Conserved roles of GATA4 and its target gene TBX2 in regulation of human cardiogenesis"

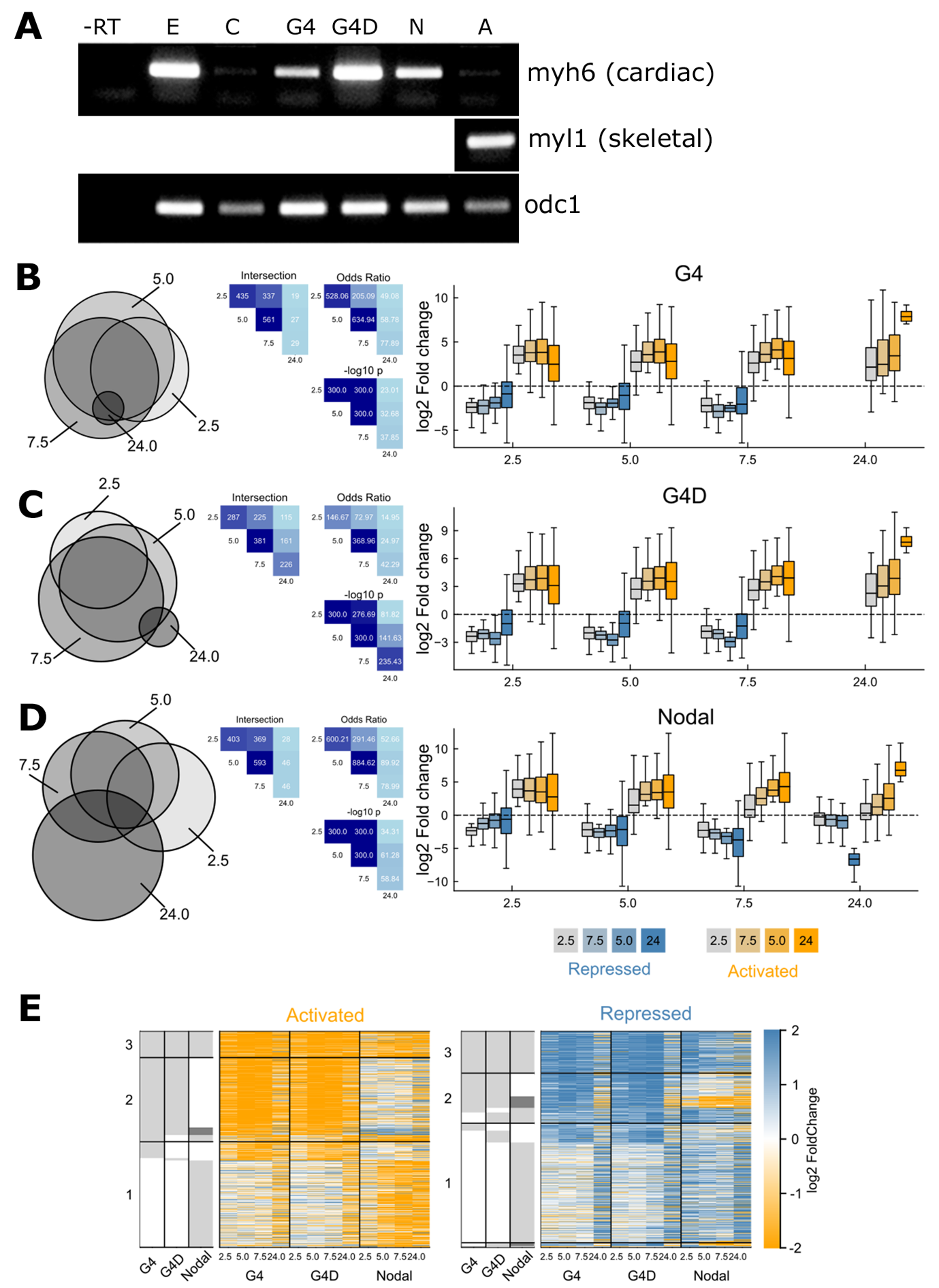

**Figure S1.** **Summary of differential expression time course for genes induced by GATA4 (G4), GATA4+DKK1 (G4D), and Nodal5 (N, Nodal) in *Xenopus* animal cap explants.** (A) Confirmation of cardiogenic conditions used in RNA-seq. RT-PCR showing expression of cardiac marker *myh6* in animal cap explants exposed to cardiac inducing conditions (G4, G4D and Nodal) and *myl1* when skeletal muscle is induced by a low dose of Activin protein. *odc1* is used as a loading control. Samples were collected when sibling embryos reached st. 34. E- sibling embryo. C- control animal caps. (B-D)**.** Intersection between differentially expressed genes between timepoints for G4 (B), G4D (C), Nodal (D). For each condition, left provides Venn diagram of intersection between differentially expressed genes at each timepoint; center provides intersection size, odds ratio, and right-tail Fisher exact test p-values between intersection of all pairs timepoints; right boxplots describing the log2 Fold Change of genes differentially expressed at each time point (horizontal axis 2.5, 5.0, 7.5, and 24h), at all other timepoints. We find highly significant associations between all timepoints, particular 2.5-7.5h. (E) Heatmap of log fold change of activated and repressed genes per timepoint, as in Figure 1D regardless of timepoint. Left grey heatmap indicates whether gene is differentially expressed at any timepoint.

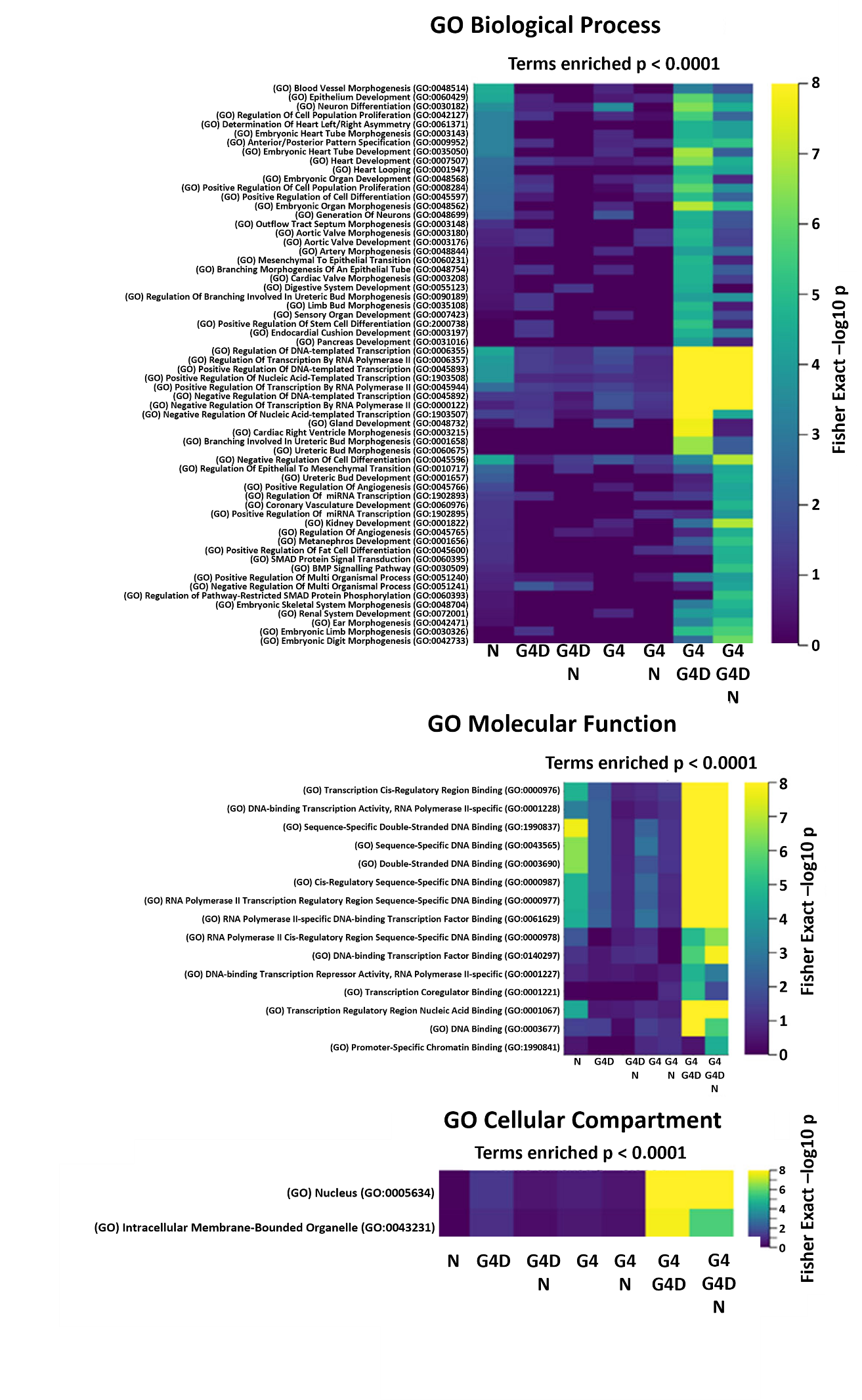

**Figure S2**. **Gene ontology enrichments for genes differentially expressed during cardiogenesis induced by GATA4 (G4), GATA4+DKK1 (G4D) and Nodal5 (N) in *Xenopus* animal cap explants.** Heatmap of gene ontology enrichments with FDR < 10-6 for sets of genes defined in upset plot in Figure 1b, heatmap as shown in Figure 1d, for Biological Process (top), Molecular Function (middle), and Cellular Compartment (bottom).

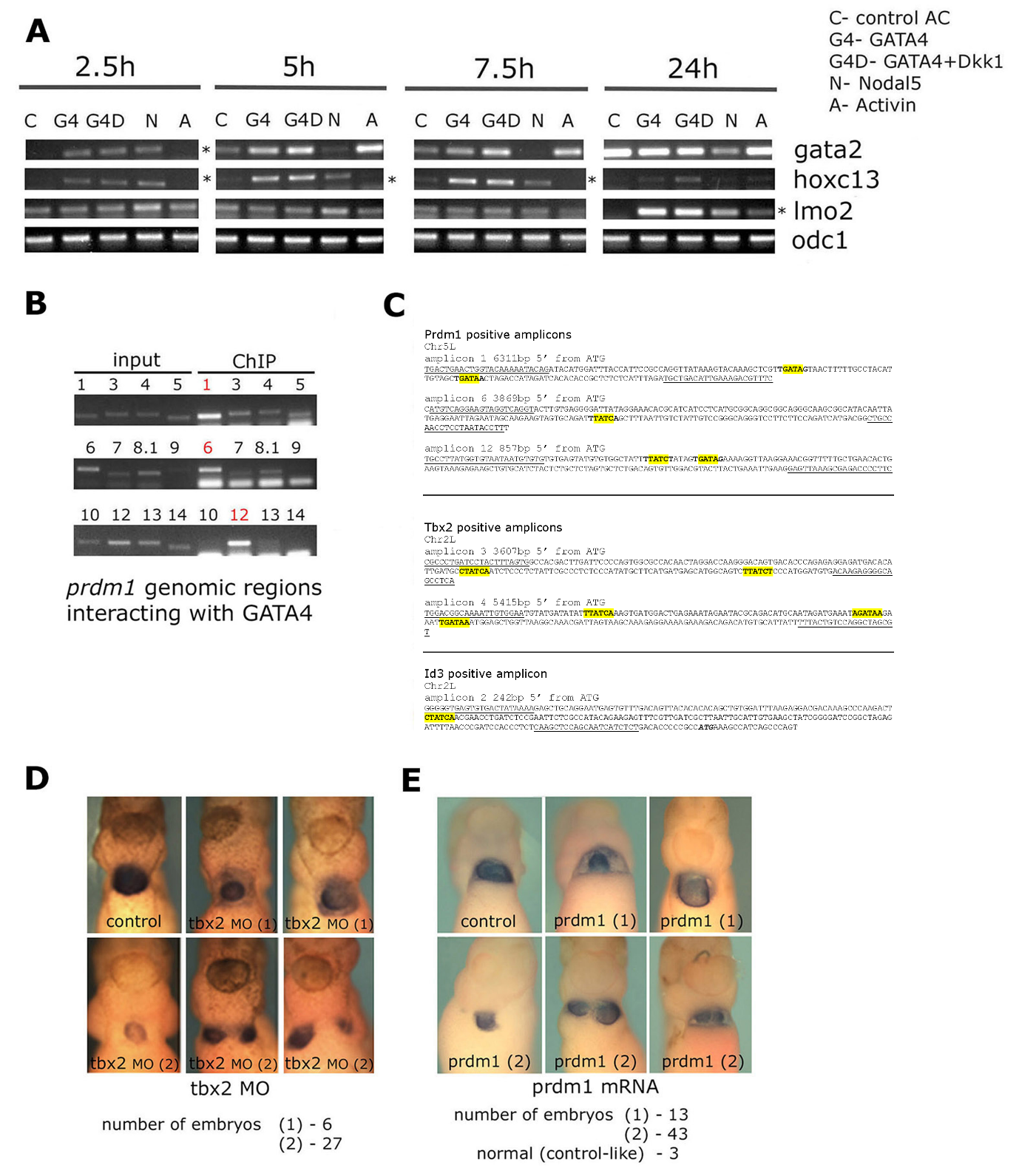
**Figure S3**. **Additional *Xenopus* DEG validation data.** (A) Validation of further DEGs by RT-PCR. *gata2* is validated at 2.5h, but not at later time points when low dose of Activin, known to induce blood, induces the expression of this haematopoietic transcription factor. *hoxc13* is validated at all time points except 24h, whereas *lmo2* is only validated at 24h, when the level of expression induced by low dose of Activin is clearly lower than the level of expression induced by Nodal5, as well as G4 and G4D. *odc1* is used as a loading control. The same *odc1* data is shown in all panels where the same cDNA samples were used. (B) Three out of twelve tested *prdm1* genomic regions interact with GATA4. These are indicated by red numbers (1, 6, 12) in gel analysis of PCRs. The region 12 is the same as region 2 in Fig. 2F. (C) Sequences of *prdm1*, *tbx2* and *id3* ChIP-positive amplicons. Sequence used to derive PCR primers is underlined. Translation start ATG was used as starting point for numbering the sequences 5’ from it. GATA sites are highlighted in yellow. (D, E) Additional images for in vivo analysis of *tbx2* and *prdm1* function. Ventral views of the head and heart region of st. 36/7 *Xenopus* *laevis* embryos processed for *myl7* whole mount in situ hybridisation to visualise the heart. Embryos injected with *tbx2* MO or *prdm1* mRNA were classified in 2 groups: (1) with milder phenotype (looping defects) and (2) with more severe phenotype (smaller linear heart tube, cardia bifida). The number of embryos in each group is indicated. Control embryos were uninjected. Anterior to the top.

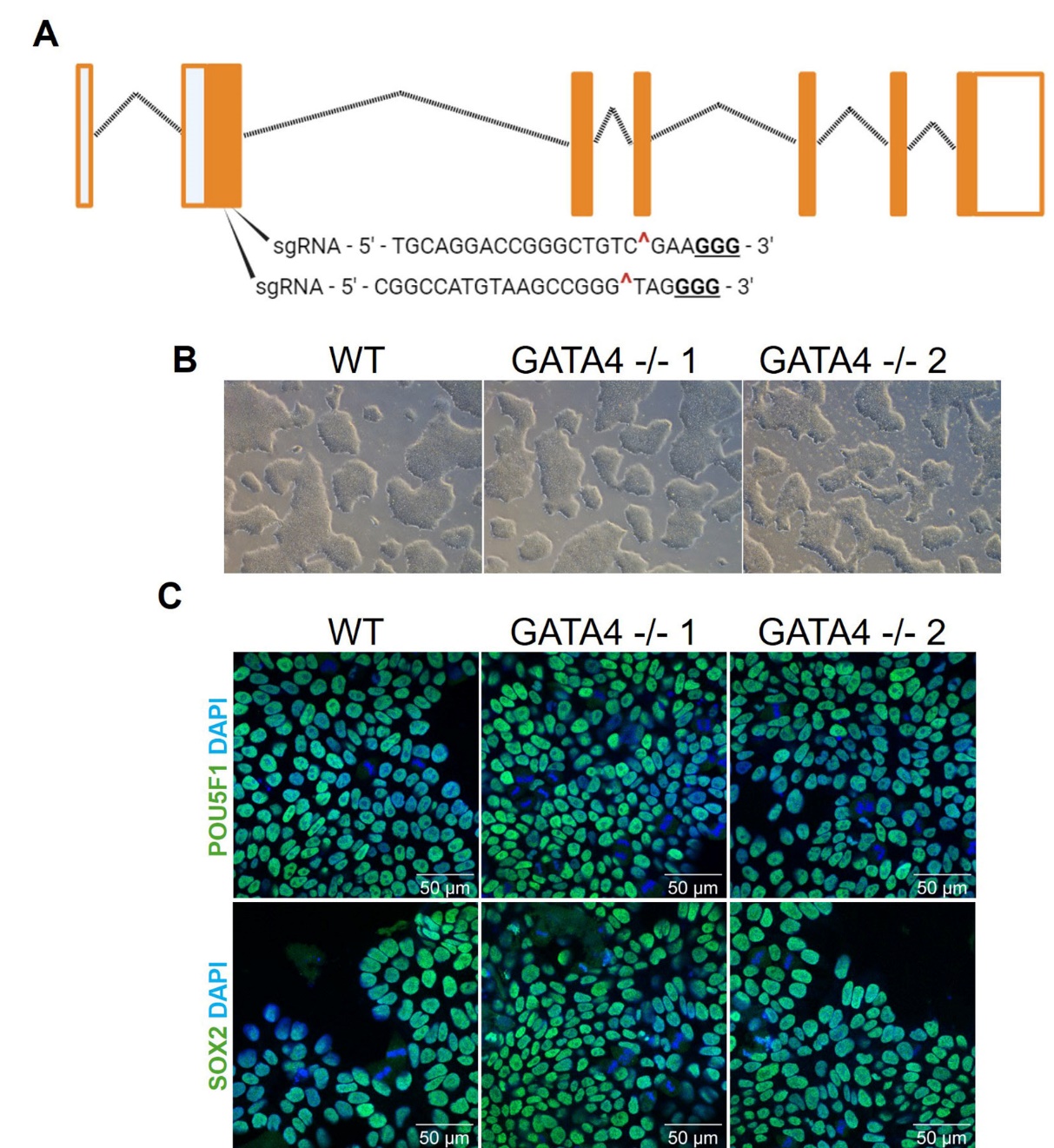

**Figure S4. Generation and pluripotency assessment of the of GATA4 -/- iPSC lines**. (A) An overview of the human GATA4 gene locus. Exons are represented as boxes; coding regions are filled versus non-coding unfilled regions. Introns are represented by a dotted line. The CRISPR guides to edit the gene were targeted towards the end of the first coding exon and are labelled as sgRNA. The PAM for each is shown in bold and underlined. The cut site is indicated by a ‘^’. (B) Brightfield 4x images of WT and GATA4 null iPS cell colonies in culture. (C) Immunofluorescence for POU5F1 (green), and SOX2 (green) in WT and GATA4 null iPS cell colonies. DAPI (blue) was used to counterstain nuclei. Two independent lines with identical trans-heterozygous deletions (not shown) were created. In all figures samples labelled GATA4^-/-^ were derived from the GATA4^-/-^1 line.

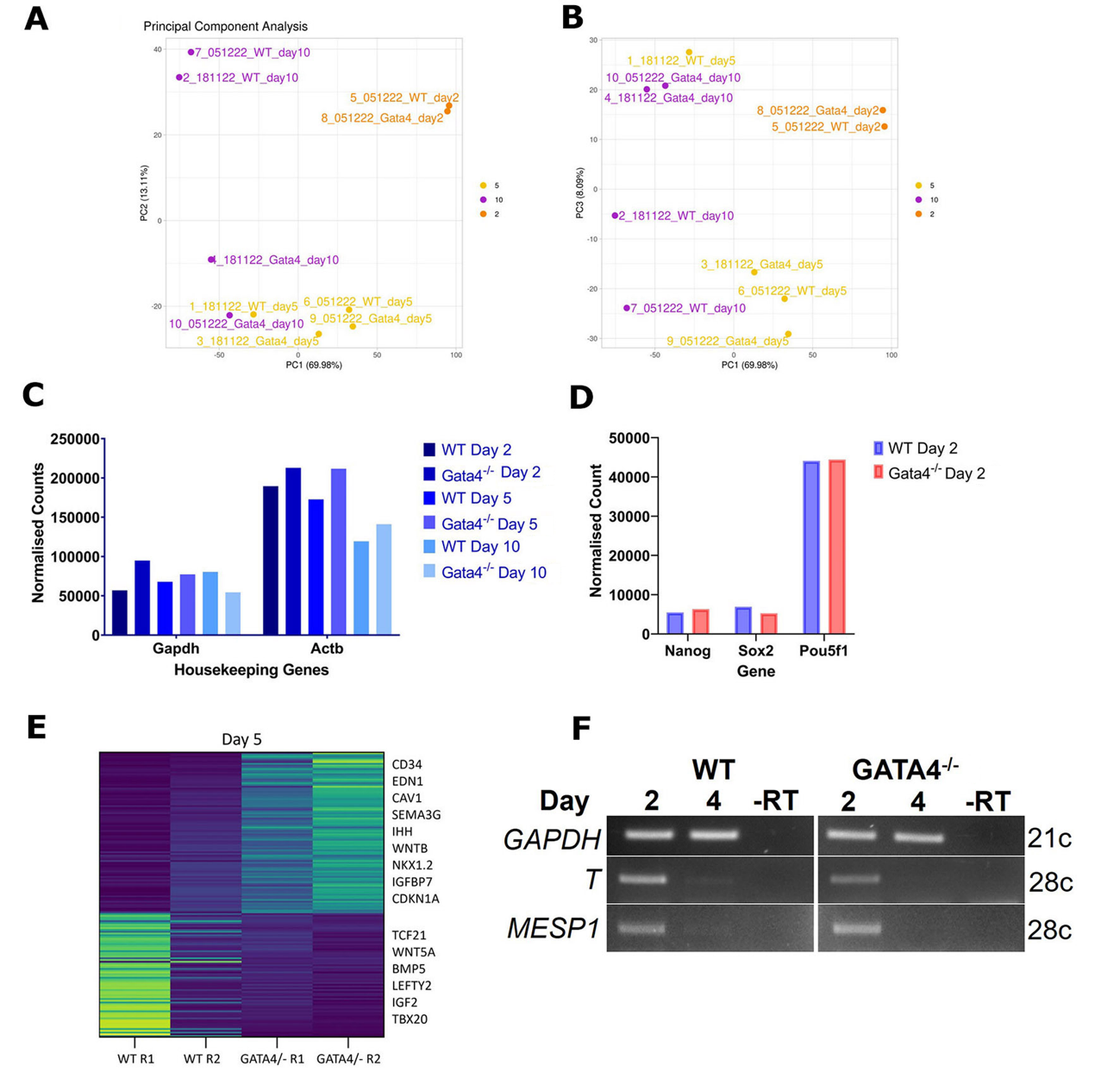

**Figure S5.** **Quality control of GATA4 -/- iPSC-CM RNA-seq data**. (A-B) Principal component plots for WT and GATA4-/- samples at days 2 (n = 1), 5, and 10 of differentiation, n = 2. (C) The mean average of the normalised counts for housekeeping genes; *GAPDH* and *ACTB* are show for each time point. (D) A plot of normalised counts for WT and GATA4-/- samples at day 2 for pluripotency genes *NANOG*, *SOX2*, and *POU5F1,* n =1. (E) Heatmap of all genes found to be differentially expressed (DEGs) by a log2FC of 2 or more in the GATA4 -/- line at day 5, two repeats (R1, R2). Example genes are indicated on the right. (F) RT-PCR analysis showing comparable expression of *T* and *MESP1* in WT and GATA4-/- cells at days 2 and 4 of differentiation. *GAPDH* has been used as a loading control (n = 2).

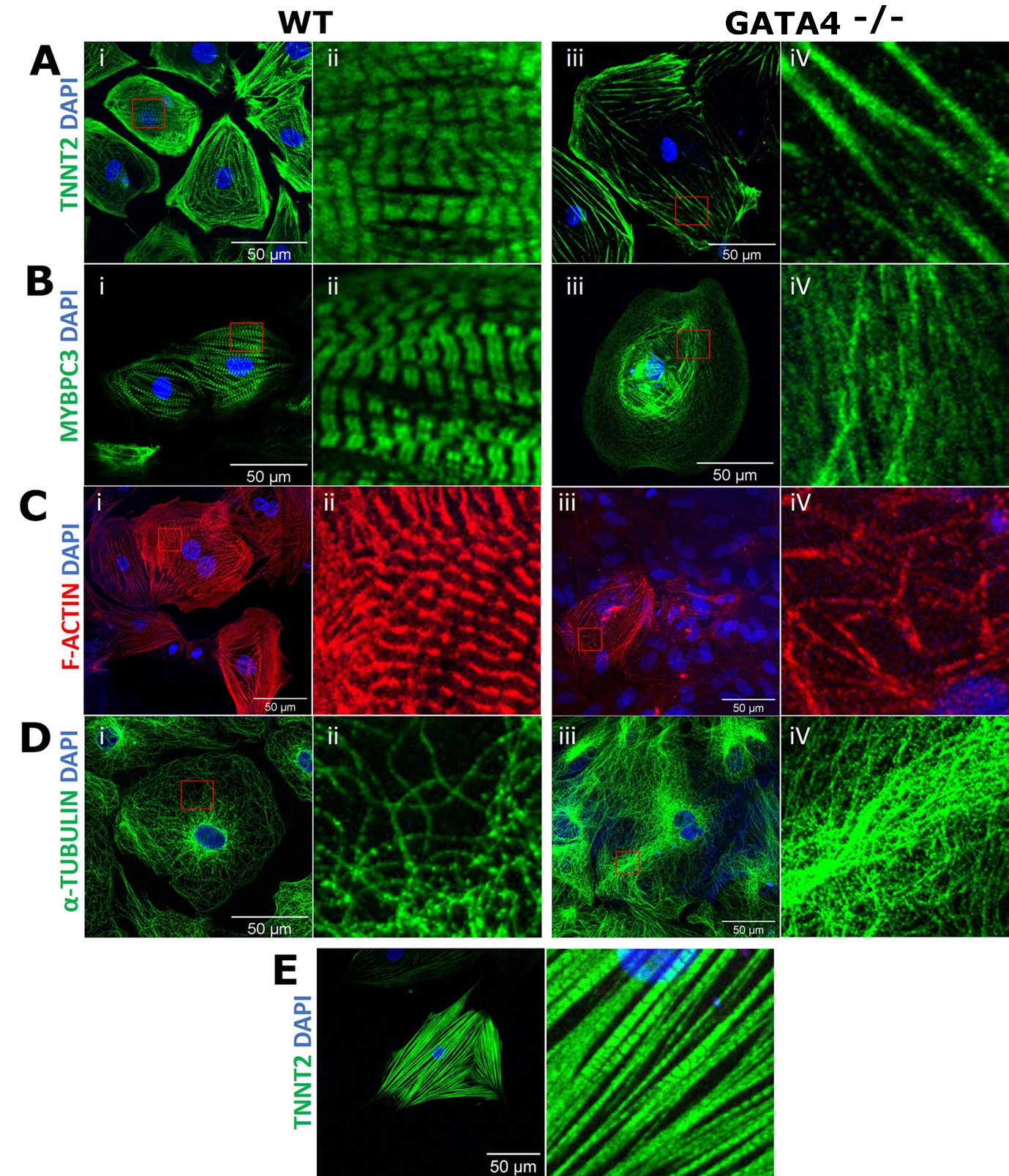

**Figure S6. Expression of sarcomeric components in WT and GATA4 null cells.**

IF staining was carried out on WT and GATA4-/- lines at day 32 and images taken at 40x with an oil immersion lens. Panels (A-C) show staining for sarcomeric components TNNT2, MYBPC3 and Filamentous Actin (F-ACTIN). Panel (D) shows staining for non-sarcomeric cytoskeletal protein α-TUBULIN. Images labelled ii and iv show cropped images enlarged so that any striations present can be more easily observed. In panel (E), a GATA4-/- cell with some striations is shown. Among the GATA4-/- cells analysed, this one had the mildest phenotype.

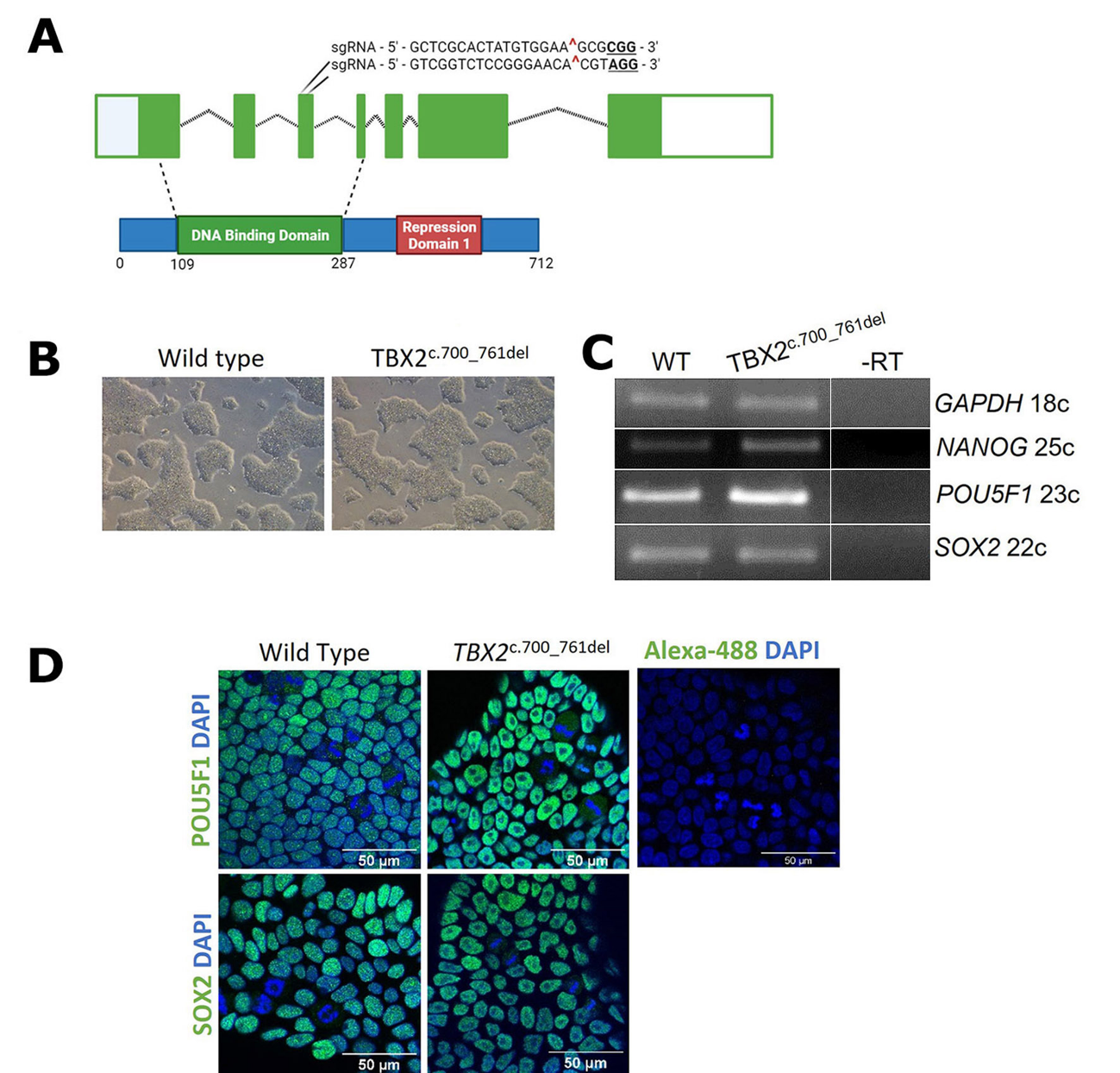
**Figure S7. Editing of the *TBX2* gene locus and pluripotency control of the mutant line.** (A) A representation of the *TBX2* gene locus with exons shown as boxes. Coding regions are filled, and non-coding regions are empty. Introns are represented by a dotted line. The CRISPR guides used for editing were targeted to exon 3. The resulting mutation leads to the entirety of this exon being skipped. See Fig. S9 for more details. This is mapped onto the protein representation below to indicate the impact of this on the DNA binding domain. (B) Brightfield images showing WT and TBX2 c.700_761del iPSC colonies. (C) Expression of pluripotency factors NANOG, POU5F1, and SOX2 analysed by RT-PCR in WT and TBX2 mutant iPSCs. GAPDH is used as a normalisation control. (D) Immunofluorescent images of WT and TBX2 mutant iPSCs stained for POU5F1 and SOX2 (green), and counterstained with DAPI (blue). Taken at 63x with a confocal microscope.

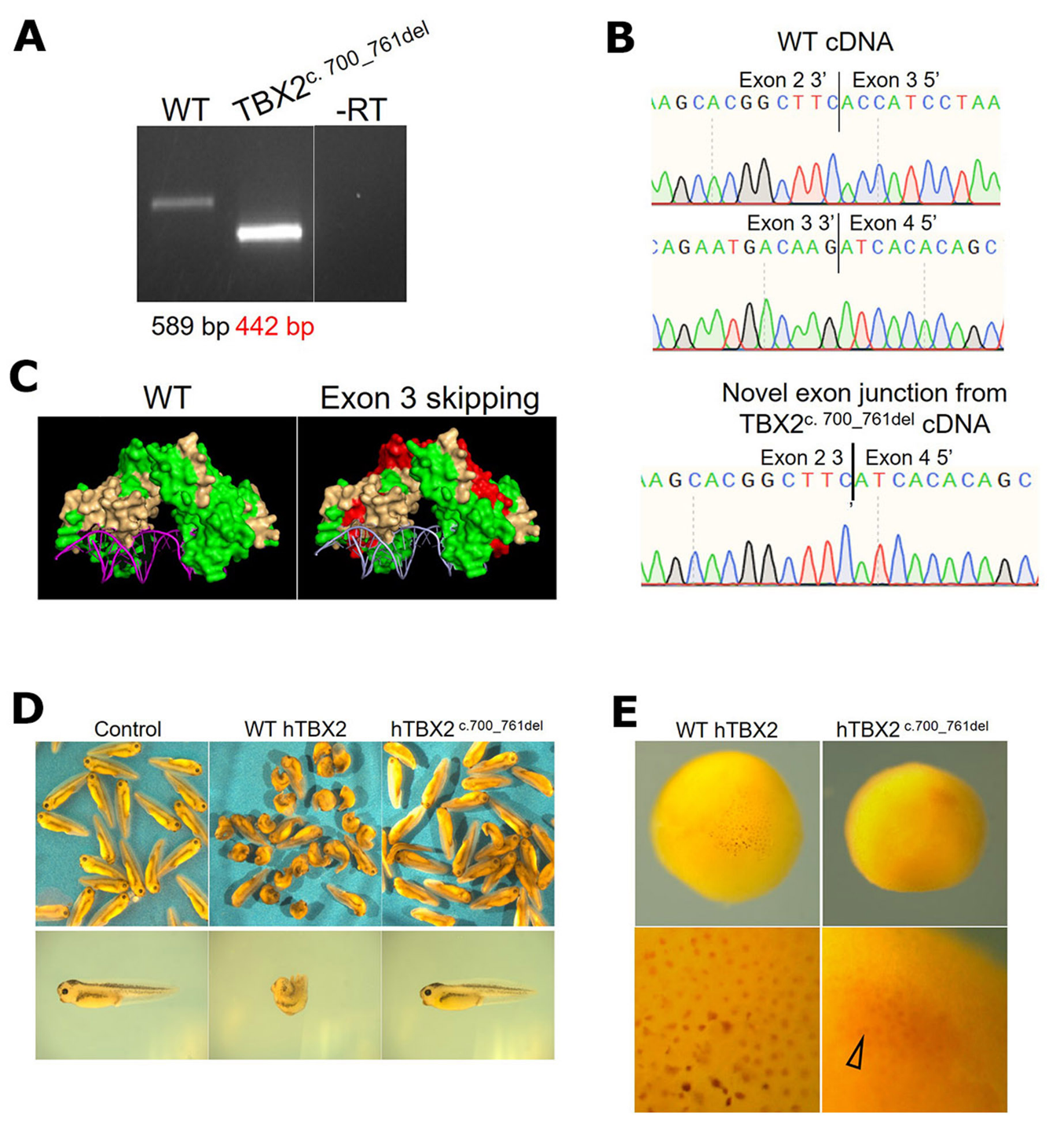

**Figure S8. The outcome of the TBX2 mutation.** (A) RT-PCR analysis of day 10 samples shows based on the cDNA produced that the mRNA transcribed from the mutated gene locus misses ~150 bp. (B) Sanger sequencing trace files confirm 147bp deletion. The areas covering the 3’ and 5’ exon junctions from WT cDNA are shown alongside the new exon junction present in the TBX2 mutant that skips exon 3 (147 bp). (C) TBX3 is the closest relative of TBX2 with a 3D structure available. Here, the TBX3 DBD (PDB ID 1H6F) is shown binding to DNA as a homodimer. The regions absent in the TBX2 c.700_761del mutant line are shown in red. (D) Brightfield images showing stage 37/8 *Xenopus laevis* embryos, injected uniformly at the 2-4 cell stage with a WT h (human)TBX2 (n = 93) or hTBX2 c.700_761del (n = 97) capped mRNA, in comparison to un-injected controls (n = 101). Single pictures displaying representative example embryos from each treatment group are shown below. The embryos injected with WT TBX2 mRNA show strong gastrulation defects, but those injected with mutant TBX2 mRNA are indistinguishable from controls. (I) Immunohistochemical staining in stage 9 embryos using an anti-HA antibody to detect HA tagged WT and mutant hTBX2 proteins. The WT but not mutant protein is readily detected in the nuclei. Taken together, the data shown here and in Fig. 6A indicate that the deletion of aa. 700-761 removes part of DNA binding domain and leads to destabilisation of TBX2 protein, resulting in a functional null mutation, at least when expressed in *Xenopus* embryos. Indirect assessment of TBX2 mutant protein in *Xenopus* embryos was undertaken as commercially available antibodies against human TBX2 showed low sensitivity.

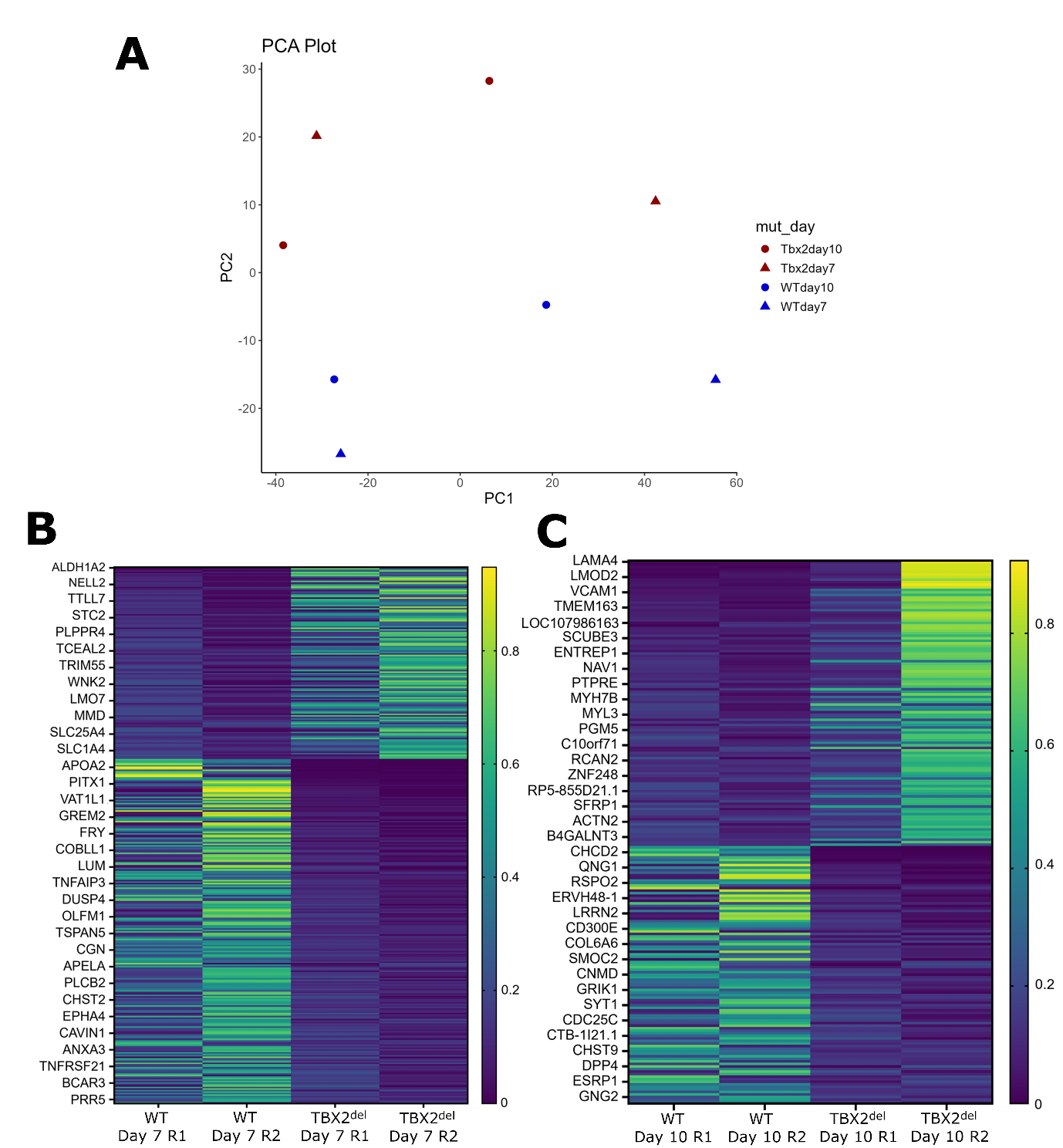

**Figure S9.** **Quality control of TBX2 -/- iPSC-CM RNA-seq data and selected up- and down-regulated genes**. (A) PCA plot showing WT vs TBX2 mutant samples at day 7 and day 10 of differentiation. (B) A heatmap showing DEGs at day 7 in the TBX2 mutant line and (C) at day 10. Whilst the upregulated genes and GO terms are more prominent (p values in the 10^-11^-10^-20^ range; Fig. 6, Supplementary Table 2), the downregulated genes and GO terms (p values in the 10^-5^-10^-9^  range; Supplementary Table 2), include ECM genes such as COL6A6, RSPO2 and SMOC2 (panel C) and GO terms such as ‘ECM’, ‘extracellular region’ and ‘collagen-containing extracellular matrix’ ( Supplementary Table 2).

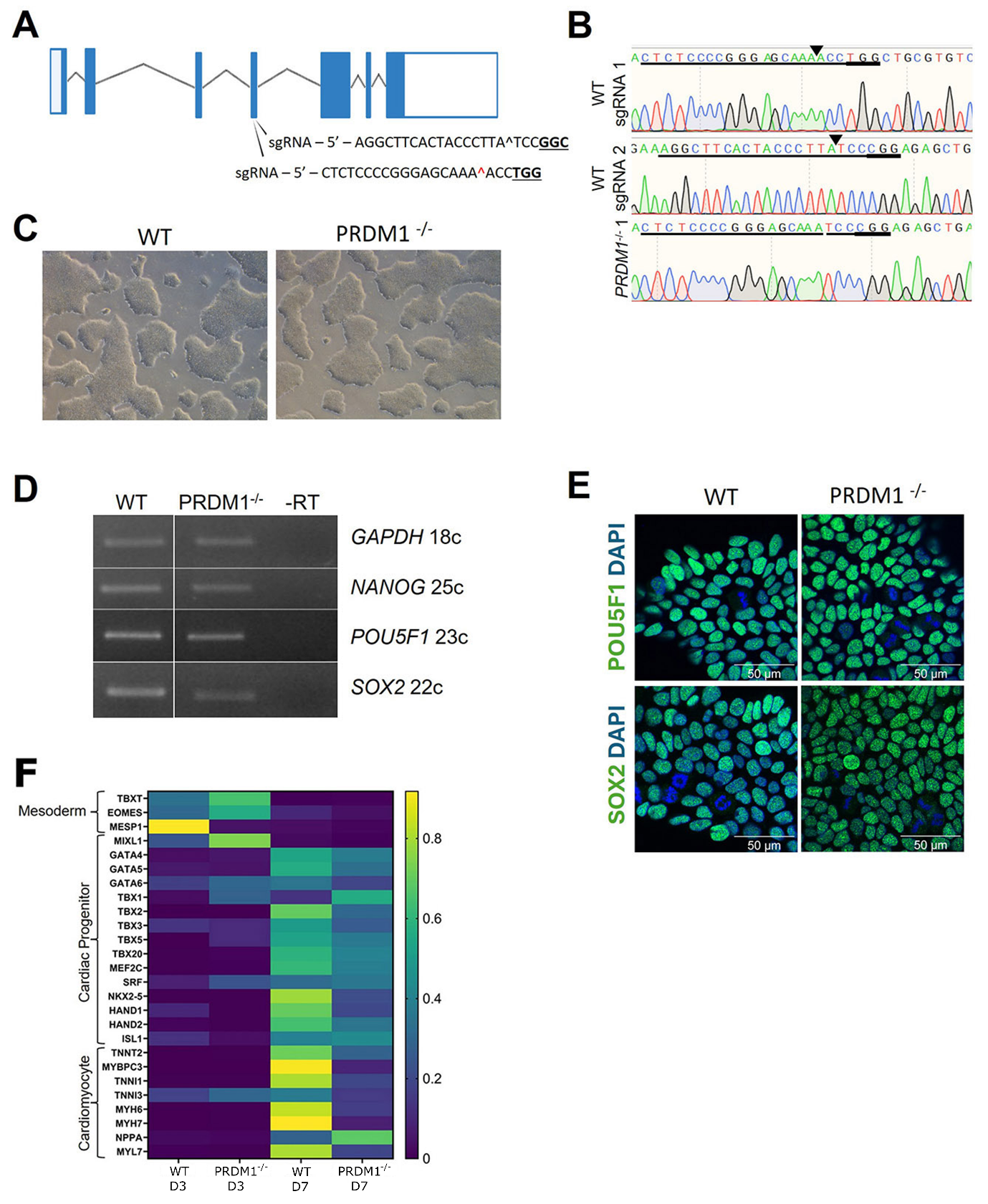

**Figure S10. Generation and pluripotency assessment of the PRDM1 -/- iPSC line**. (A) The PRDM1 gene locus with exons depicted as boxes and coding regions of these as filled boxes. Introns are represented as dotted lines. The guide RNAs used to knockout PRDM1 were targeted to exon 4 as this is shared amongst all PRDM1 isoforms. The PAM sequence is shown in bold and underlined. The cut site is indicated by ‘^’. (B) Sanger sequencing trace files showing the locus before and after editing. Guide RNAs have been underlined with PAM sites underlined in bold. Cut sites indicated with a black arrow. (C) Brightfield images showing WT and PRDM1-/- iPS cell colonies. (D) RT-PCR showing the expression of NANOG, POU5F1, and SOX2 expression in WT and PRDM1-/- 1 iPSCs. GAPDH was used as a normalisation control. (E) Immunofluorescence images showing POU5F1 (green), and SOX2 (green) expression in WT and PRDM1-/- cells with DAPI (blue) used to counterstain nuclei. This demonstrates localisation of these TFs to the nucleus in non-dividing cells. (F) A heatmap generated from RNA-seq data showing the expression of mesoderm, cardiac progenitor and cardiomyocyte markers as days 3 and 7 of differentiation in the WT and PRDM1 -/- line.
